# Antigen experience relaxes the organisational structure of the T cell receptor repertoire

**DOI:** 10.1101/2021.10.26.465870

**Authors:** Michal Mark, Shlomit Reich-Zeliger, Erez Greenstein, Dan Reshef, Asaf Madi, Benny Chain, Nir Friedman

## Abstract

The creation and evolution of the T cell receptor repertoire within an individual combines stochastic and deterministic processes. We systematically examine the structure of the repertoire in different T cell subsets in young, adult and LCMV infected mice, from the perspective of variable gene usage, nucleotide sequences and amino acid motifs. Young individuals share a high level of organization, especially in the frequency distribution of variable genes and amino acid motifs. In adult mice, this structure relaxes and is replaced by idiotypic evolution of the effector and regulatory repertoire. The repertoire of CD4+ regulatory T cells was more similar to naïve cells in young mice, but became more similar to effectors with age. Finally, we observed a dramatic restructuring of the repertoire following infection with LCMV. We hypothesize that the stochastic process of recombination and thymic selection initially impose a strong structure to the repertoire, which gradually relaxes following asynchronous responses to different antigens during life.

## Introduction

The ability to sustain effective T cell immunity relies on a diverse αβ heterodimeric T cell receptor (TCR) repertoire generated by the stochastic variable, diversity and joining (VDJ) recombination mechanism(Kohler et al., 2005). This diverse repertoire is shaped over time by recombination biases (Qi et al., 2014)(Snook et al., 2018), thymic and extra- thymic selection(Kohler et al., 2005) (Qi et al., 2014) (Kavazović et al., 2018), selective migration and antigen-driven clonal expansion. The encounter with cognate peptide-MHC complex (pMHC) also drives the differentiation of the T cell. For example, the strength of TCR stimulation can skew differentiation of memory versus effector T cells(Snook et al., 2018) (Kavazović et al., 2018) and CD4+ regulatory (Treg) versus effector/memory CD4+ cells(Lee et al., 2012) (Stritesky et al., 2012) linking TCR specificity to phenoytpe and function. The aim of this study is to document the influence of these diverse processes on the underlying structure and organization of the TCR repertoire, determined at a global level.

Several previous studies have used deep sequencing to explore the TCR repertoire in different T cell subsets. For example, significant changes can be found between the repertoires of CD4+ and CD8+ cells, presumably reflecting selection by different classes of MHC peptide complexes(Li et al., 2016)^-^(Gulwani-Akolkar et al., 1995). Similarly, the repertoire differences found between CD4+ Treg and conventional CD4+ cells(Pacholczyk et al., 2006),(Wang et al., 2010) are presumed to be shaped by their recognition of self or foreign peptides. However, the processes driving repertoire diversification are probabilistic, rather than deterministic. As a result, identical TCR sequences can be found in multiple subsets, and can even be shared between CD4+ and CD8+ populations(Wang et al., 2010).

In young individuals, the majority of the T cell compartment is made up of naïve cells, and the repertoire is presumably shaped largely by stochastic recombination and thymic selection. However, as individuals age their immune system responds to an increasing number of foreign antigens, derived principally from microbial, allergen or altered-self (e.g. neoantigen) exposure. This drives a relative shift towards the memory/effector phenotype(Arnold et al., 2011), accompanied by increased clonal expansion. Interestingly, exposure to antigen in different individuals can drive both convergent and divergent repertoire evolution (Heather et al., 2016),(Pogorelyy et al., 2018). At the repertoire level clonal expansion results in a gradual decrease in overall repertoire diversity(Jörg J. et al., 2015) (Britanova et al., 2014). The CD4+ T cell repertoire diversity is more preserved with age in the bone marrow compared to the spleen(Shifrut et al., 2013), which may relate to the role of the bone marrow microenvironment in preservation of memory T cells (Di Rosa and Pabst, 2005),(Baliu-Piqué et al., 2018). The Treg repertoire also changes with age, as production of thymic “natural “ Treg drops significantly, and are replaced by a high proportion of Tregs with active effector/memory phenotype(Smigiel et al., 2014),(Thiault et al., 2015).

In this study, we combine multi-parameter fluorescence-activated cell sorting with high-throughput-next generation sequencing to undertake a comprehensive high resolution analysis of the αβ TCR repertoire of various T cell compartments in young and adult mice, comparing CD4+ and CD8+ T cells of naïve, central memory, effector and Tregs, from the spleen and bone marrow. We illustrate the impact of strong antigen exposure on the global properties of the repertoire by analyzing the changes that follow infection with lymphocytic choriomeningitis virus. We quantify the global parameters of the repertoire at different levels of dimensionality, spanning variable gene frequencies, amino acid motif frequencies and at the level of individual nucleotide sequences. We explore different ways to visualize the structure and order which underlies the superficially diverse and chaotic collections of different DNA and protein sequences which constitute the T cell repertoire. Finally, we interpret our observations from the perspective of the probabilistic, but not chaotic processes which determine the development and evolution of the TCR repertoire. We hypothesize that these processes operating on millions of T cells impose a strong overall structure to the repertoire. This structure relaxes as a result of divergent responses to antigen exposure in different individuals.

## Results

### A quantitative description of the TCR repertoire

We collected CD4+ and CD8+ T cells of naïve, central memory, effector and Tregs, from the spleen and bone marrow of 12 and 52 week old mice (summarized in Fig 1A). Representative flow cytometry plots showing the phenotypic markers, the gating strategy and relative purity of the populations obtained are shown in supplementary (SI Fig 1A-B). We appreciate that our antibody panel does not fully capture the complexity of the T cell compartment, and that more extensive panels would be required to fully differentiate between all the known sub-compartments. However, for the purpose of this high level analysis, we simplify the nomenclature, and refer to the sorted populations as naïve, Treg, central memory and effector. After RNA extraction, we amplified the TCR repertoire using a previously published experimental pipeline which incorporates unique molecular identifiers (UMI) for each cDNA molecule to correct for PCR bias and sequencing error, allowing a robust and quantitative annotation of each sequence in terms of V gene, J gene, CDR3 sequence and frequency (Oakes et al., 2017),(Uddin et al., 2019).

**Figure 1.**
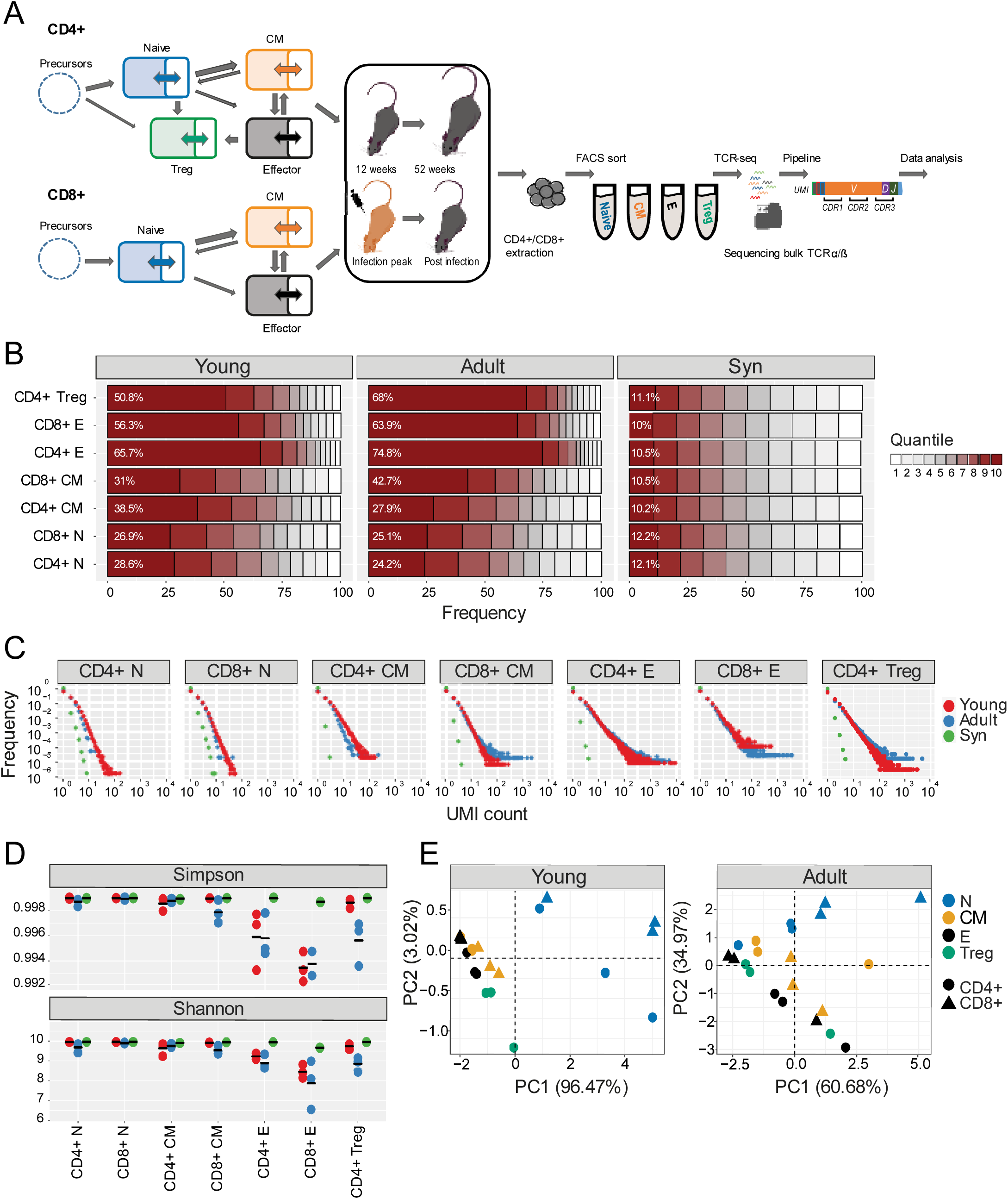
Clonal expansion and diversity of the TCRβ repertoire in different subsets of young and adult mice. **(A)** Summary of T cell compartments and pipeline for cell isolation and TCR repertoire sequencing and analysis. (**B**) The TCRs in each repertoire were ranked according to frequency, and the proportion within each decile is illustrated (low abundance sequences in white, ranging to high abundance sequences in dark red). The percentage of the distribution represented by the top decile is shown in white text. (**C)** The sequence abundance distribution in each compartment. The plots show the proportion of the repertoire (y-axis) made up of TCR sequences observed once, twice etc. (x-axis). Repertoires from young mice are shown with red dots, repertoires from older mice in blue dots and synthetic repertories in green. **(D)** Simpson and Shannon scores of equal repertories size (1000 CDR3NTs) from each compartment and mouse. Colors same as panel C. Mean is shown in black lines (n=3). **(E)** PCA of the Renyi diversities of order 0, 0.25, 0.5, 1, 2, 4.

The numbers of cells and the number of TCR mRNAs (captured by the total UMI count) which were recovered varied widely between compartments and age groups. For example, both splenic CD4+ and CD8+ naïve compartment from young mice resulted in the highest average UMI count (∼415,000) while the splenic CD4+ central memory (CM) population yielded the lowest average UMI count (∼44,000). As expected, the proportion of naïve cells in both spleen and bone marrow was higher in young than adult mice, and this was balanced by an increase in memory and especially effectors in the older mice (SI Table 1). The total UMI count was strongly correlated with the number of sorted cells across compartments and tissues (SI Fig 1C). The number of α and β UMIs were also highly correlated (SI Fig 1D). Both these correlations provide additional confidence in the robustness and quantitative output of the overall pipeline.

### The clonal structure and diversity of the repertoire varies with compartment and age

We first explored the changes in the clonality and diversity of the TCR repertoire across compartments and tissues. We estimated T cell clonotype size by the number of different UMIs associated with a unique TCR, and illustrated the clonal frequency distribution of the repertoire within each population (e.g. Figs 1B and 1C for spleen; SI Fig2A and B for bone marrow). As a comparator in this, and subsequent figures, we generated a set of synthetic TCRs using SONIA, a generative probabilistic model of TCR recombination which incorporates learnt parameters of the genomic TCR recombination process, without any subsequent selective expansion(Sethna et al., 2020). This serves as a useful baseline with which to compare real repertoires, in which the products of recombination have been shaped by selection and proliferation.

As expected, the naïve repertoires were dominated by rare TCRs (observed only once or twice in a sample) and had very few expanded clonotypes (expanded clones are represented by the darkest color in panel B, and by the points to the right in panel C). The naïve repertoires were also most similar to the synthetic repertoires. In contrast, T effectors contained much larger numbers of expanded clonotypes, and this was more pronounced in CD8+ cells from the older mice. Consistent with these distributions, the Simpson index, and the Shannon index, two commonly used measures of diversity of the repertoire, were highest in naïve populations from young individuals, and progressively lower in central memory and effectors (Fig 1D, SI Fig 2C). The Simpson and Shannon indices are examples (k = 2 and k = 1, respectively) of a series of diversity measurements, which are captured by the Renyi entropy of order k, where k can run from 0 to infinity. We calculated the Renyi diversities for k = 0, 0.25, 0.5, 1, 2, 4 for each repertoire and then plotted them in two dimensions using principal component analysis (PCA; Fig 1E and SI Fig 2D). In the young mice, the repertoires of naïve, central memory, effector and T regulatory cells are clearly separated by the diversity measurements alone, with almost all the variance captured in a single dimension (reflecting very consistent differences across the entire Renyi profile). In older mice, the distinction between the populations is still observed but is less clear cut, and with greater variation between individual mice. All panels in Fig 1 show the results obtained for the TCRβ repertoires (spleen), because TCRβ repertoires are the most diverse and are more commonly studied. However, similar results were observed for the α repertoires, and the diversity of α and β repertoires was very highly correlated (SI Fig 1E).

In summary, the analysis of the repertoires of different populations captures the known decreasing diversity and increasing clonality of the naïve, central memory and effector compartments in both spleen and bone marrow and the decrease in diversity observed with age. These results build further confidence in the reliability of the repertoire sequencing and analysis pipeline.

### Differential V gene usage defines different sub-populations of T cells in young individuals

As reported previously (Ndifon et al., 2012),(Madi et al., 2014), both Vα and Vβ gene usage was non-uniform in all the repertoires examined, and also in the synthetic repertoire sequences, reflecting differential usage of V genes in the recombination process (Kohler et al., 2005)(SI Fig3A). However, the distribution of V gene usage also differed between T cell naïve subsets (young vs. adult, adult vs .synthetic, young vs. synthetic mice). The pairwise similarity between V gene distributions of different repertoires was quantified using the cosine similarity between the distributions (see Methods). We also used the Horne similarity index(Greiff et al., 2015),(Venturi et al., 2008) and found these two measures highly correlated (SI Fig3B).

A hierarchical clustered heatmap summarizes the similarity between all pairwise combinations of repertoires for TCRβ V genes (Fig 2A). In young individuals there was a clear segregation between CD4+ and CD8+ repertoires, and between naïve, central memory, effector and Treg populations. Naïve, central memory, and Tregs repertoires were most similar, while effectors were mapped to a distinct branch. In contrast, there was little distinction between spleen and bone marrow within each sub-compartment. Repertoires from the same compartment but different individuals clustered together, demonstrating that each compartment had a distinct repertoire distribution, conserved between individuals.

**Figure 2:**
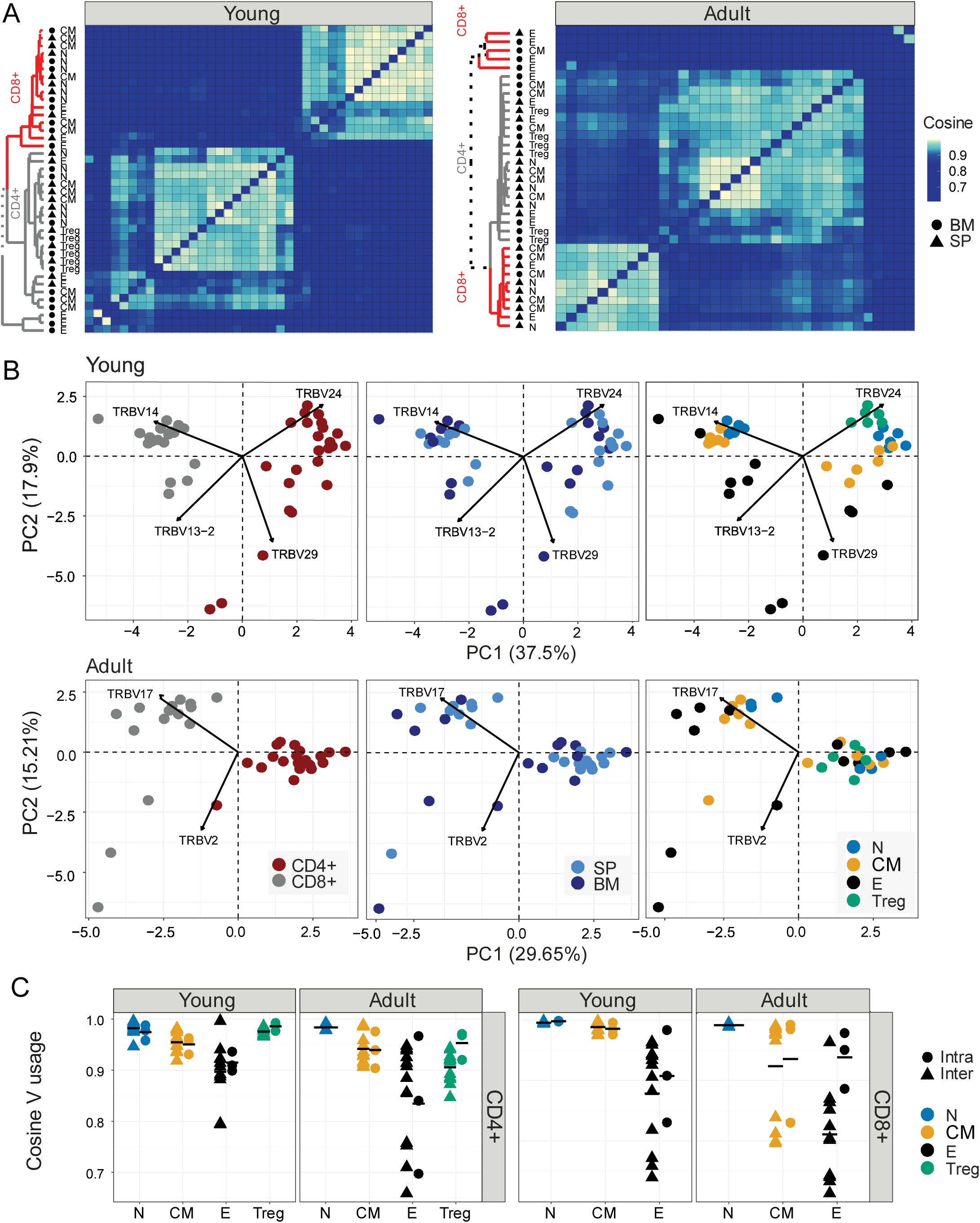
Differential V gene usage defines different sub-populations of T cells in young individuals. (**A)** Cosine similarity was calculated between all pairs of repertories in young (left) or adult (right) mice and displayed as a heatmap. Hierarchical clustering dendrograms showing the organization of the assigned at each plot, colored by CD4+ and CD8+ groups (grey and red branches respectively) and labels by compartment (text and symbol). Tissues are marked in symbols shape (SP = triangles, BM = circles). **(B)** PCA separates the Vβ usage of CD4+ and CD8+ compartments in age dependent-manner (Young in upper and Adult in lower panel). Each color represents one compartment from one mouse (e.g., CD8+ Effectors, BM, mouse 1). See legend for symbols and color code. PC1 separates between CD4+ and CD8+ classes in both age groups. PC2 divides between cell compartments in Vβ usage of young mice. The Vβ genes with the highest influence (loading) are marked with arrows. (**C)** Cosine similarity of the Vβ gene usage between individuals (circles) or within individuals (between spleen and bone marrow, triangles). T cells compartments (colored dots) are divided to CD4+ (left) and CD8+ (right) from young or adult mice. Mean is shown by horizontal black lines.

In contrast to the strong hierarchical structure observed in the TCRβ repertoires in 12-week individuals, the Vβ gene usage in repertoires from older animals was much more heterogenous. Although the distinction between CD4+ and CD8+ repertoires was mostly still retained, the sub-compartments were much more inter-mingled. The repertoires of the CD8+ effector compartments, in particular, showed little similarity between individuals.

The structure observed in the heatmap organization was further investigated by performing principal component analysis (PCA) on the pairwise similarity matrix for Vβ usage (Fig 2B) for young (top panels) and adult (bottom panels) mice. Each dot represents an individual repertoire and is colored by CD4+/CD8+ compartment (left panels), anatomical compartment (middle panels) and differentiation phenotype (right). In young mice there is a clear separation of both CD4+ and CD8+ repertoires, and of repertoires from different functional compartments. We noted that the Treg populations lie closest to the naïve, while the biggest variance is seen between effector populations. In adult mice, the separation between CD4+ and CD8+ repertoires is retained, but the distinction between functional compartments largely collapses.

In contrast to the TCRβ repertoires, the equivalent analysis for the α repertoires (SI Fig 3C and 3D) showed much less evidence of consistent structure in either heatmap or PCA. Furthermore, there was only limited correlation between the cosine similarities of α and β repertoires, especially in the older individuals (SI Fig 3E). The selective pressures which shape the repertoires of different CD4+ and CD8+ compartments therefore seem to be reflected differently in Vα and Vβ gene usage.

Since we observed that there were no systematic differences between spleen and bone marrow repertoires in terms of Vβ gene distribution, we estimated the degree of variation which could be attributed to idiosyncratic differences between mice, by comparing intra-individual (between bone marrow and spleen) differences with inter-individual differences (Fig 2C). The plots illustrate a clear hierarchy of variance, with naïve repertoires being closest to each other, followed by central memory and Tregs, and with effector repertoires showing the greatest divergence. CD8+ repertoires (right panels) showed greater divergence (smaller similarity indices) that CD4+ repertoires, and the adult repertoires showed greater variance than young. Interestingly the intra-individual variation was in general very similar to the inter-individual variation, the only exception being the effector CD8+ repertoires in the older animals. Thus, the high variance seen especially between effector T cell repertoires seems to be an intrinsic property of these repertoires, observed even between different compartments from the same individual. This high variance was not simply a reflection of the different sizes of the different compartments since different sized synthetic repertoires were very similar to each other (SI Fig3 F-G).

### T cell sub-compartments defined by nucleotide sequence sharing patterns

The TCR V gene distributions analyzed above create a simplified abstraction of individual repertoires, and TCR repertoires can also be considered as a hyperdimensional feature space defined by the millions of individual nucleotides which constitute each repertoire. In order to identify structure within this space, we first visualized the qualitative patterns of sharing between CD4+ and CD8+ sub-compartments, using circus plots (Fig 3A). This analysis, which included only sequences shared by at least two compartments, reveals a distinctive pattern of sharing which is conserved between individuals, and is age specific. In young individuals, CD4+ and CD8+ splenic naïve and CD8+ central memory repertoires contribute the highest proportion of shared sequences (blue [0.21-0.26, 0.28-0.39, CD4+ and CD8+, respectively] and orange [0.29-0.39])circus arc lengths. Naïve repertoires from adult mice contribute a much smaller proportion (0.004-0.03, 0.03-0.12, CD4+ and CD8+ respectively) of sharing with other repertoires, and CD4+ (0.307-0.313, 0.12-0.23) and CD8+ (0.195-0.375,0.11-0.23) effectors sequences now make up the largest proportion of shared sequences (blue, black, and grey, circus arc lengths, in SP and BM respectively). Interestingly, high levels of overlap (0.172-0.307) are observed between splenic CD4+ Treg and CD4+ naïve repertoires, while in adult mice, Tregs become more similar to CD4+ effector cells (0.159-0.290), this observation is investigated in more detail below. Nucleotide sequence sharing between T cell compartments was explored in more detail using the cosine similarity index to quantify pairwise inter-repertoire TCR sharing between compartments. Because the similarity between repertoires of different individuals at nucleotide level is very low, we first analyzed each mouse separately. However, visual inspection suggested the patterns obtained for all three mice was very similar, especially for the younger individuals, and this was confirmed by quantitative comparisons of the similarity indices between the different mice (SI Fig 4A). A representative heat map of all pairwise comparisons for a single mouse is shown in Fig 3B (TCRβ) and SI Fig 4B (TCRα), and the similarity matrix is visualized in two dimension using multidimensional scaling in Fig 3C (TCRβ) and SI Fig 4C (TCRα). In young mice a hierarchical structure was observed, with naïve and Treg repertoires clustered together, and effector and central memory repertoires for CD4+ and CD8+ T cells forming distinct clusters. In older individuals, this structure is perturbed. CD4+ and CD8+ repertoires remain distinct, but Tregs now cluster independently of naïve, and are closest to CD4+ effector repertoires. As was the case for V gene similarities, there was modest correlation between TCRα and β similarities, especially in the older individuals (SI Fig 4D). The synthetic repertoires show very little sharing or structure, consistent with clonal expansion being driven by selective forces which operate subsequent to recombination (Fig3 B-C, SI Fig4 B-C, right subplots).

**Figure 3:**
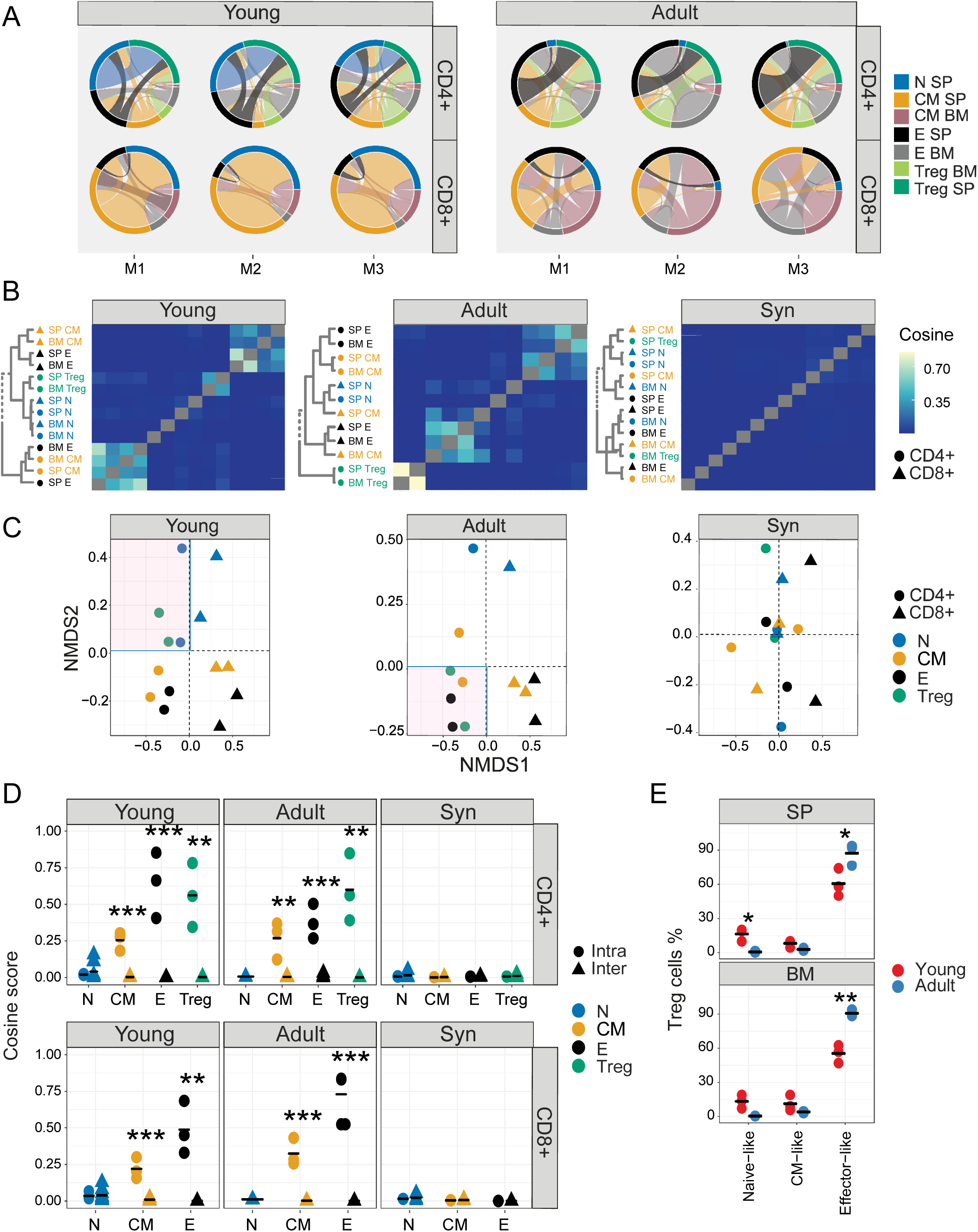
Differential sharing of T cell CDR3 nucleotide sequences defines different sub-populations of T cells that change with age. **(A)** Each circus plot represents a single mouse CD4+ or CD8+ compartment (upper and lower panel, respectively). Circus sharing levels illustrate the number of clones shared between two compartments (band widths), and the proportion of shared clones attributed to each compartment (circus arcs). Only sequences shared by at least two compartments were included in the analysis. **(B)** CDR3βNT sequences pairwise cosine similarity from representative young, adult or synthetic (“Syn”) mouse repertoires. Correlation levels are represented by color (high=light blue, low= dark blue). Hierarchical clustering dendrograms for all T cell compartments are plotted to the left of each heatmap (CD4=circle, CD8= triangles), in color and text. **(C)** The similarity matrices shown as heatmaps in B are represented in two dimensions by NMDS. **(D)** Cosine similarity between CDR3NTβ chains across (triangles) and within individuals (between spleen and bone marrow, circles). T cells compartments (colored dots) are divided to CD4+ (left) and CD8+ (right) from young (upper), adult (middle) or synthetic (“Syn”, lower) mice repertories. Mean is shown by horizontal black lines. **(E)** The surface phenotype of Foxp3+ Tregs. The plot shows the percentage of Foxp3 positive cells (Treg): CD44- CD62L+ (naive-like), CD44+CD62L+ (CM-like) and CD44+CD62L- (effector-like). Mean is shown by horizontal black lines. Each data point represents one mouse. Significant differences between age groups or intra and inter individuals are denoted by asterisks (P-values: ^*^ <0.05, ^**^ < 0.01, ^***^ < 0.001, with FDR correction t-test).

The nucleotide similarity hierarchy is illustrated in more detail for all three mice for selected compartments in Fig 3D and SI Fig 4E. Inter-individual similarity index at nucleotide level is very low in all compartments. The cosine similarity between spleen and bone marrow (i.e. intra-individual) is lowest for naïve repertoires, reflecting high diversity and limited clonal expansion. It increases for central memory repertoires, and is highest for T effectors and Tregs, reflecting lower diversity and increased clonal expansion. Strikingly, the overall intra-individual hierarchy observed is reversed compared to V region usage. Treg repertoires were more similar to themselves than to other repertoires, but more similar to CD4+ effector repertoires in older than in younger mice (SI Fig 4E). The shift from a naïve-like to an effector-like Treg observed from the perspective of repertoire sharing was also observed in phenotype, with a higher proportion of FoxP3+ CD62L+ CD44- naïve Tregs in young animals, and a higher proportion of Foxp3+ CD62L-CD44+ effector-like Tregs in the older animals (Fig 3E).

### T cell compartments defined by differential frequency of amino acid motifs

The extreme hyper-dimensionality of the sequence space dominates individual patterns of clonal diversity and expansion, and limits the recognition of conserved repertoire organization. We and others (Thomas et al., 2014),(Glanville et al., 2017) have shown that short patterns of sequential amino acids (k-mers) can play a key role in determining specificity, and offer one way to reduce the dimensionality of the repertoire while reflecting the complexity of antigen recognition. We therefore counted the presence of sequential amino acid triplets (dimensionality 3^20) or 7-mers (dimensionality 7^20) in each repertoire. To further reduce the dimensionality of the feature space, we removed rarely used features as described in detail in the Methods.

The distribution of triplet and 7-mers frequencies are represented in two dimensions by the first two components of a PCA. The k-mer distributions separated CD4+ and CD8+ TCRβ repertoires in both young and older mice (SI Fig 5A and SI Fig 5B). In the younger repertoires, conserved distinct patterns of k-mer frequency were also evident between the naïve, Treg, central memory and effector CD4+ sub-compartments (Fig 4A and SI Fig5C), with Tregs lying close to the naïve, and central memory repertoires lying between naïve and effectors. This clear hierarchy became much more relaxed in the older individuals. Within the CD8+ compartment, central memory and naïve cells cluster together, and the overall pattern is driven by a high variance of the CD8+ effectors, which diverge from each other both within and between individuals. A similar qualitatiative pattern was seen for TCRα triplets and 7-mers, although the distinction between naïve and central memory was evident in both CD4+ and CD8+ compartments (SI Figs 6A and SI Figs 6B). The intra-individual and inter-individual cosine similarities are summarized in Figs 4B (triplets) and SI Fig 6C (7-mers).

**Figure 4:**
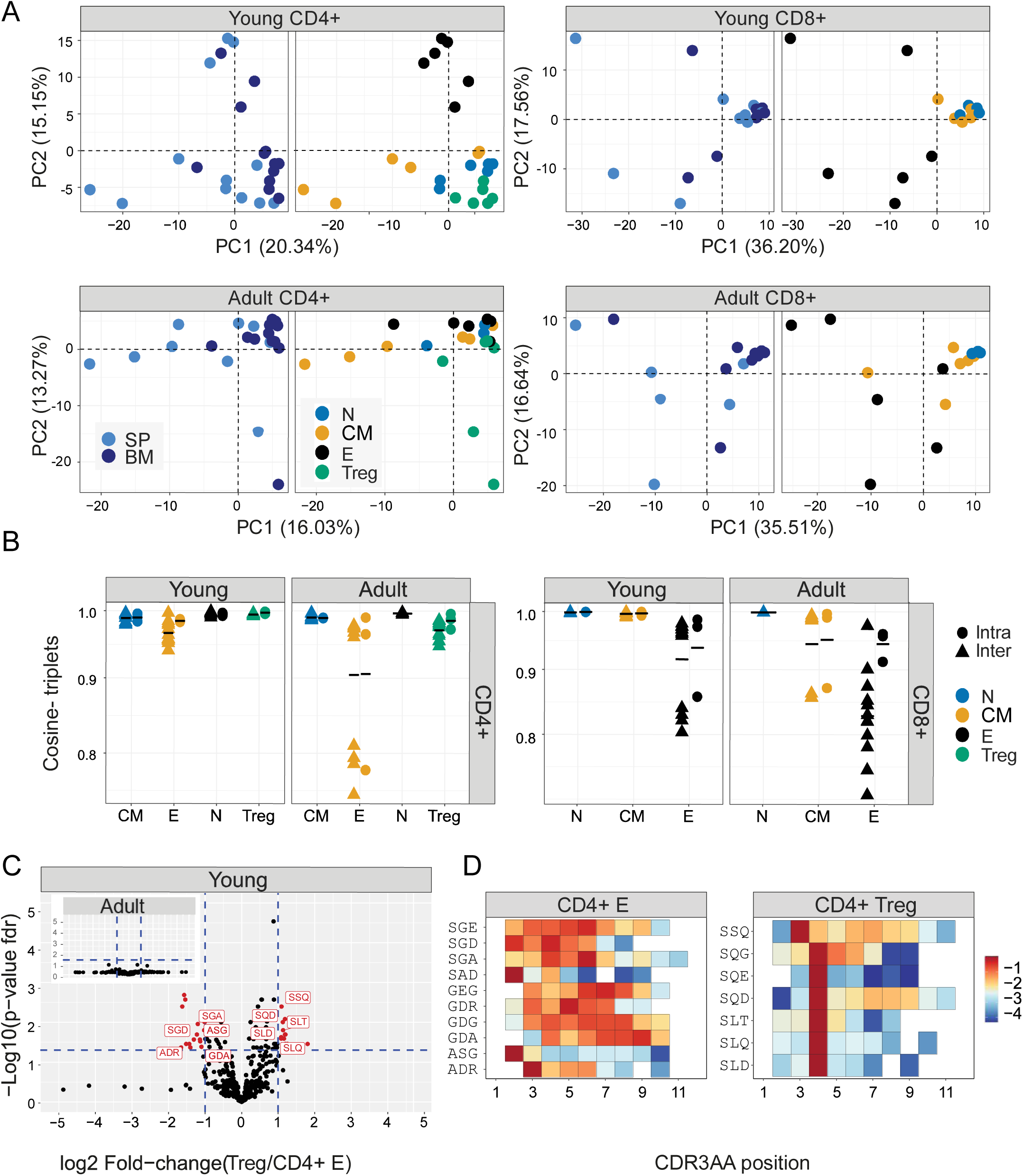
CD4+ T cells compartments distinct top CDR3βAA triplets motifs, alter with age. Top sequential triplets are selected by the mean frequency of each motif across all compartments and mice **(A)** CDR3βAA triplets PCA analysis of CD4+(left) or CD8+(right) from young (upper) or adult (lower) mouse (e.g., CD8+ effectors, BM, mouse 1). **(B)** Cosine similarity of the top (350) CDR3βAA triples between individuals (circles) or within individuals (between spleen and bone marrow, triangles). T cells compartments colored dots) are divided to CD4+ (left) or CD8+ (right) from young or adult mice. Mean is shown by horizontal black lines. **(C)** Treg and CD4+ effector differentially expressed triplets are found in young but not adult mice. Each dot represents a single triplet (- top or all 8000 triplets in red or black dots, respectively). P-value (t-test) was calculated for each triplet across six samples (three mice and 2 tissues) of CD4+ Treg and CD4+ effector cells. The y-axis shows FDR-adjusted p-values. The x-axis shows the log 2-fold-change, calculated between Treg and CD4+ effector mean triplets or motifs frequency across compartments (6 samples in each). Significance thresholds are marked in blue lines: (1) at y=1.3 (equivalent to p-value of 0.05) and x=±1 (denoting a total fold-change of 2). Representative triplets above both thresholds are labeled with red text and dots. **(D)** Significantly expressed triplets positioned in various positions along the CDR3AA sequences. Triplets overexpressed in CD4+ Treg are frequently located in position 4 of the CDR3AA’s. (3-9). Triplets overexpressed in CD4 effector can be located mainly in position 2-3 or further along the CDR3AA sequences. The color represents the log10 frequency of each aligned triplet.

Interestingly, and similarly to what we observed in V gene distributions, the inter-individual similarities were only consistently larger than the intra-individual similarities for the CD8+ effectors.

We examined in more detail the differential usage of amino acid motifs between Treg and T effectors (Fig 4C, SI Fig 7A). In younger repertoires ten triplet motifs were over-represented in the CD4+ effector repertoires, and seven in the Treg repertoires. In the older repertoires there was little evidence of differential motif use between these compartments (see insets). Almost all the differentially-represented triplets began with a serine (Fig 4D). The triplet motifs over-represented in the Treg repertoires were found almost exclusively at positions 3/4 of the CDR3 suggesting they may be acting as a surrogate for selective V genes; however the triplets over-represented in the T effectors were more broadly distributed across the CDR3 (Fig 4D). The 7-mers over-represented in the CD4+ T effectors were predominantly found associated with a single V gene. In contrast, the 7-mers over-represented in Treg repertoires were more broadly distributed (SI Fig 7B). Overall, while V gene usage plays a part in the amino acid motif distribution profiles, selection independent of V gene is clearly at work.

### Combined feature sets which incorporate V gene, nucleotide and amino acid motif distributions can discriminate T cell sub-compartments in young individuals, but these differences weaken with age

The results presented in figs 2-5 illustrate how different feature sets offer quite distinct perspectives on the organisation of the TCR repertoire. We concatenated all the intra-individual intra-compartment, and the inter-individual intra-compartment (spleen only) cosine similarity values from all feature sets (i.e. V gene, nucleotide, triplet and 7-mers) into a single vector, and displayed a representation of this feature space in two dimensions by applying PCA (Fig 5). In this representation, the distinction between CD4+ and CD8+ repertoires were lost. However, in the repertoires of young individuals, a clear organization separating naïve, central memory, effector and Tregs is now evident. In contrast, in adult mice the organization is mostly lost, and the populations are inter-mingled.

**Figure 5:**
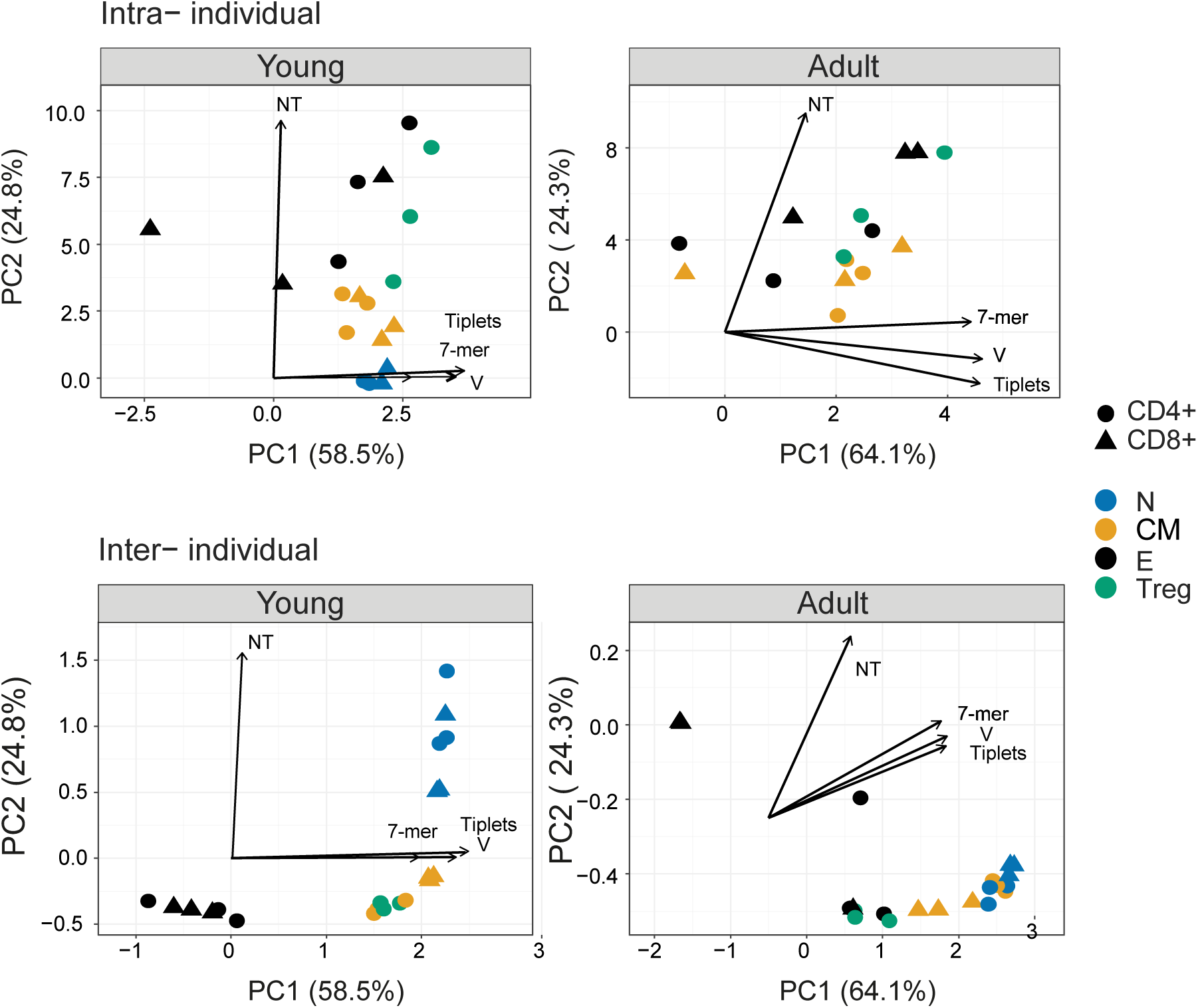
PCA analysis of the combined TCRβ features separates between T cells states of young mice. Cosine similarity calculated for the Vβ usage, CDR3βNT, top CDR3βAA triplets and 7-mers motifs between T cells compartments within individuals (between spleen and bone marrow, for example: Treg BM and Treg SP from young mouse 1) or splenic compartments across mice (for example: Treg SP mouse 1 and Treg SP mouse 2). The TCRβ measurement with the highest influence is marked with arrows (V = TRBV, NT = CDR3NT, Triplets = CDR3AA triplets, 7-mer = CDR3AA 7-mers).

### The impact of LCMV infection on repertoire organization

Finally, we examined the effect of exposure to a strong immunogenic stimulus on the organization of the immune repertoires (Fig 6A). We infected C57BL/6 mice with LCMV, which drives a strong but self-limiting infection associated with a well-characterized immune response in this strain. The cosine similarity for each compartment between mice, as well as between repertoires of young and older uninfected individuals is shown for V gene, CDR3 nucleotide and amino acid triplets (Fig 6B). Infection drives a strong decrease in similarity (increase in diversity) between naïve and memory repertoires of different mice, especially evident in the V gene and triplet distributions. In the effector population, in contrast, infection drove exactly the reverse process, increasing similarity between infected individuals, and thus counteracting the normal decrease in similarity which is observed between effector repertoires of different individuals. In this case, therefore, infection is driving convergence of the effector repertoires. The increased diversity of naïve and memory compartments is seen in both CD4+ and CD8+ populations, while the decreased diversity of the effector compartment is particularly evident in the CD8+ population. The impact of infection is strongest at 8 days post-infection, when the host response is maximal (Murali-Krishna et al., 1998),(Slifka et al., 1997)and partially returns to baseline by 40 days post infection.

**Figure 6:**
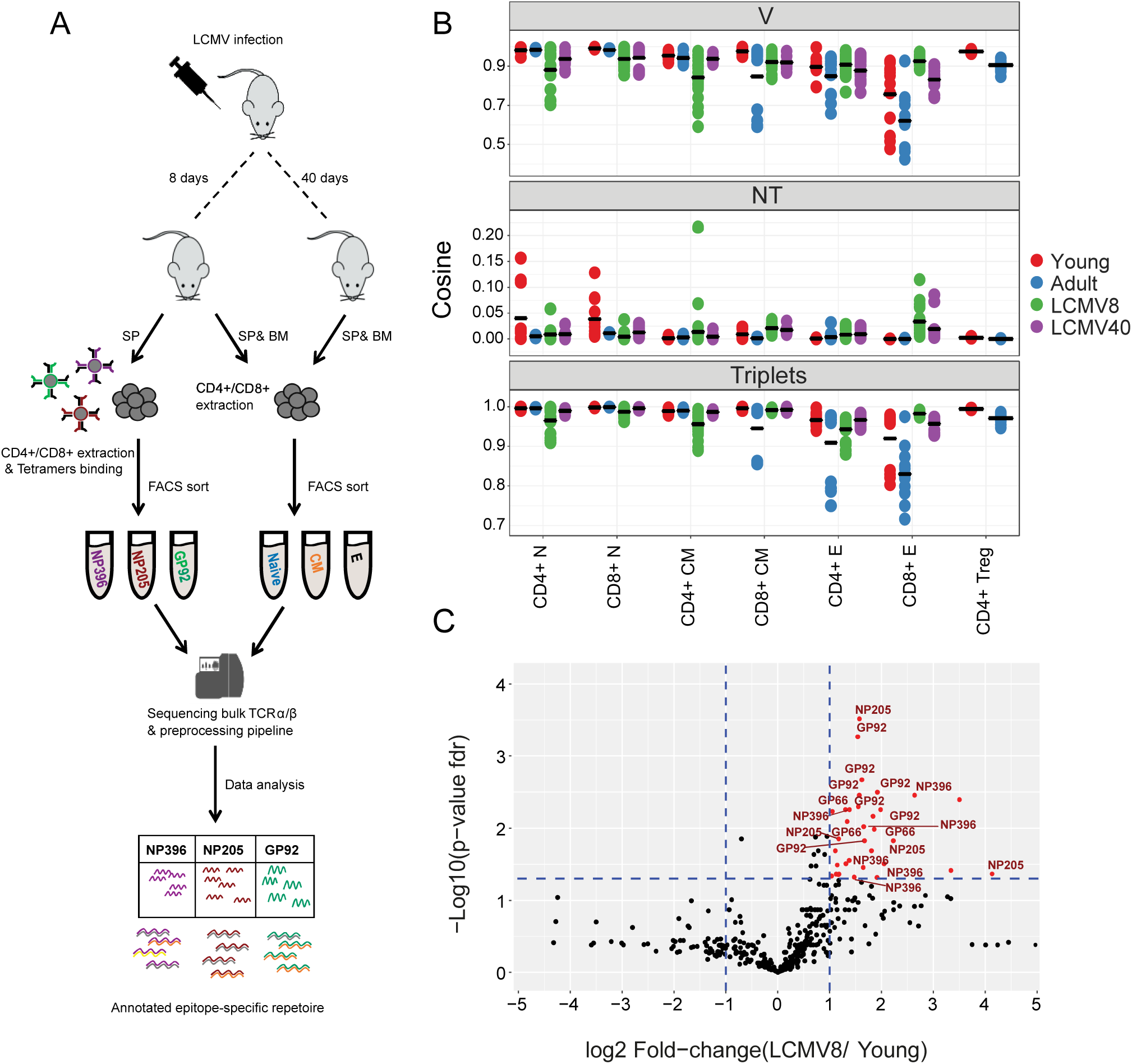
T cells compartments of LCMV infected mice express distinct top amino acid triplets of β chain TCR repertoire. **(A)** Summary of the LCMV induced T cell compartments and epitope –specific cells isolation for TCR repertoire sequencing and analysis. **(B)** Cosine similarity index of TRBV genes, CDR3βNT and top CDR3βAA 3-mers (top 350) motifs calculated between tissues and individuals. Colored dots reflect the mice groups (red = young, blue = adult, green/purple = mice after 8 and 40 days of acute LCMV infection, respectively). Mean is shown by horizontal black lines. **(C)** CD8+ effector differentially expressed triplets are found after 8 days of LCMV infection, and not in the young healthy mice. Each dot represents a single top triplet. P-value (t-test) was calculated for each triplet across six –eight samples (three- four mice and 2 tissues) of CD8+ effectors from young and LCMV infected mice. The y-axis shows FDR-adjusted p-values. The x-axis shows the log 2-fold-change, calculated between mean triplets from young and LCMV infected mice (6-8 samples in each). Significance thresholds are marked in blue lines: at y=1.3 (equivalent to p-value of 0.05) and x=±1 (denoting a total fold-change of 2). Representative triplets above both thresholds are labeled with red text and dots. Significantly enriched triplets that are labeled in red text are found in the epitope specific full CDR3βAA sequences (NP396, NP205, and GP92). 36 significantly expressed triplets are found, among them, 30 triplets are also found annotated to the epitope–specific sequences (83%).

In order to understand better the convergence observed between the effector populations of infected mice, we analysed triplet usage in the CD8+ effectors of LCMV infected versus uninfected individuals. A number of triplet motifs were highly enriched in the repertoires of the LCMV infected mice (Fig 6C, sequences in SI Table 2). Many of these triplets were also observed in the TCRs of a population of T cells isolated from the infected spleens by sorting on the LCMV peptides NP396-404(H-2D^b^), NP205-212(H-2K^b^) and GP92-101(H-2D^b^) (Fig 6A).

## Discussion

The adaptive immune system, uniquely among vertebrate physiological systems, uses a family of receptors which are not encoded in the germline, but are created de novo in each individual by a stochastic process of imprecise DNA recombination. A fundamental task for immunologists is to understand how this stochasticity and associated inter-individual heterogeneity can nevertheless result in a robust and regulated response to a enormous diversity of antigens in most individuals of a population. In this study we explore the balance between stochasticity and heterogeneity on the one hand, and order and consistency on the other. We systematically analyze the TCR repertoire of different functional and anatomical compartments of the adaptive immune system, sampled from young (3 month) and adult (12 month) mice. From this perspective, we consider the immune system as evolving in a multi-dimensional selective space. The dimensions (selective pressures) include thymic selection, peripheral differentiation (along the naïve- memory-effector axis), migration (spleen – bone marrow) and aging (illustrated in Fig 7). We document the effects of these selective processes on different features of the repertoire, which span the range from the full hyper-dimensionality of individual nucleic acid sequences (>10^8^ per mouse) through the enumeration of amino acid motifs (a few hundred), to the frequency of different V genes (20). We focus the analysis on quantitative measurements of similarity between repertoires, which reflects both convergent and divergent evolution of the repertoire. A recent study has reported systematic sequencing of TCR repertoire of different human T cell subsets, but the focus of their analysis was on the biochemical characteristics of the TCR(Kasatskaya et al., 2020) .

**Figure 7:**
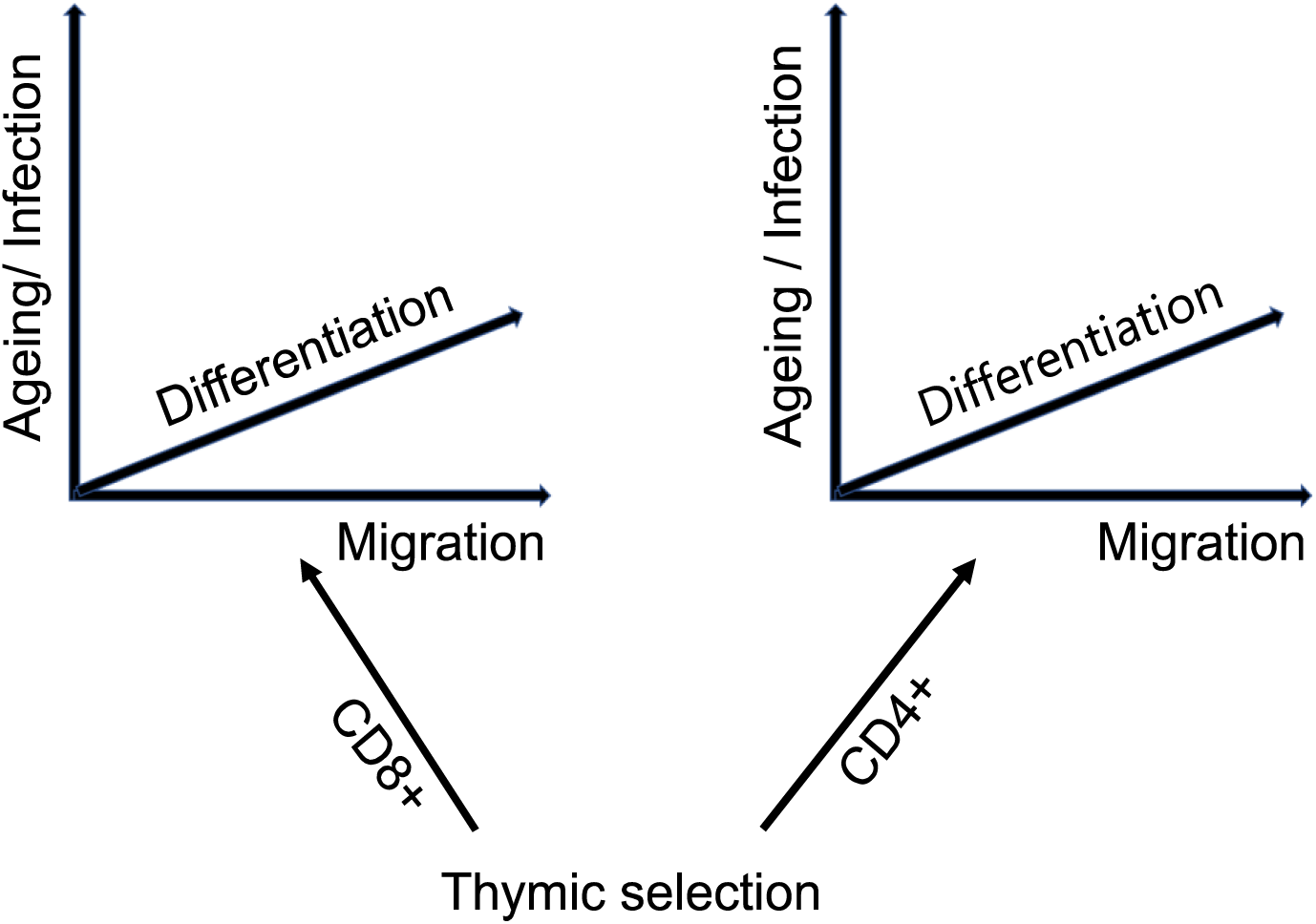
The TCR repertoire is considered as evolving in four dimensions, captured by the diagram above.

In the younger mice, the analysis of similarity revealed clear evidence of order, with a hierarchical structure of similarity between the different functional compartments. The most consistent feature was the clear separation between CD4+ and CD8+ repertoires, which was evident in all feature sets explored, in both TCRα and TCRβ repertoires, and presumably reflects the MHC/peptide selection process which operates in the thymus. Notably, however, the selection operates on a complex multi-feature construct, since no one feature (V gene, amino acid motif, or even individual CDR3 nucleotide sequence) could distinguish individual CD4+ from CD8+ TCRs. Within CD4+ or CD8+ compartments, the similarity from the perspective of V gene or amino acid motif frequency distributions was highest between naïve repertoires, with progressively decreasing similarity for memory and effector repertoires. Remarkably, this increasing heterogeneity was observed both between matched compartments of different mice and between the same compartment sampled in bone marrow and spleen. We hypothesise that this diversity is an intrinsic feature of the differentiation process shown in Fig 7, driven by clonal expansion in response to continuous exposure to a diverse set of self and non-self antigens. These selective forces must operate on the TCRα/β heterodimer, since the two genes are co-expressed as a single structure at the cell surface. However, the selection seems to operate rather independently on the α and β sequences, since the patterns of inter-repertoire sharing observed for α and β are only loosely correlated. Vβ genes are much more informative than Vα genes in terms of distinguishing functional compartments.

The tension between randomness and directed evolution is most evident when comparing the analysis of V gene frequencies and individual CDR3 nucleotide sequences. Similarity in V gene usage is greatest in naïve, and decreases progressively in central memory and effector repertoires. In contrast, similarity in CDR3 frequencies is lowest in naïve, because of the extreme diversity of this compartment, and increases progressively in central memory and effector repertoires. The combination of recombination and selection therefore impose a rigid pattern of V gene usage, which nevertheless encompasses an enormous diversity of TCR sequences. Memory and effector differentiation, presumably in response to antigen, drive some convergent evolution of the clonal repertoire, reflected by increasing similarity of nucleotide sequence repertoires, but paradoxically increasingly disturbing the rigid pattern of V gene usage.

In the older mice, elements of the structure remain, but aging and the much longer exposure to the antigenic environment significantly loosen the initial rigid structure evident in V gene and amino acid motif frequency. CD4+ and CD8+ repertoires, for example, remain clearly distinct in all feature sets. However, the clear segregation between naïve, central memory and effector repertoires becomes blurred, and the overall pattern of similarity is increasingly driven by the idiosyncratic effector repertoires which differ both at V gene and at amino acid motif level. The Treg population show a distinctive distribution of similarities. In both young and adult mice, the Treg repertoires are more similar to themselves than to any other compartment, confirming the distinct nature of the Treg repertoire, which has been hypothesized to arise from exposure to a distinct set of antigens (Wyss et al., 2016),(Bolotin et al., 2017). However, the Treg repertoires are more similar to naïve repertoires in the younger individuals, but become more similar to effector repertoires with age. The switch from a naïve-like to a more effector-like repertoire, which is also observed at a phenotypic level by increased expression of CD44 and decreased expression of CD62L may reflect a life-long gradual recruitment of induced Tregs to the original natural Treg population emerging from the thymus(Darrigues et al., 2018). The switch of regulatory T cells to a more effector phenotype might also represent a weakening of regulatory activity, and hence be linked to the increase in autoimmunity associated with age.

The response to environmental antigens drives many of the differentiation and age-associated changes which we describe. Since the mice are housed in specific pathogen free conditions, and are not germ-free, this may include a variety of microbial antigens present in the environment. However, although the mice are co-housed, the individual antigen exposure may be heterogenous and asynchronous. We therefore investigated the impact of exposure to a strong synchronous exogenous antigenic stimulus, by infecting the mice with LCMV, which produces a strong but self-limiting infection in the C57Bl/6 strain. The immune response to this virus has been studied extensively(Zhou et al., 2012), and is known to involve strong systemic clonal expansion by both CD4+ and CD8+ T cells. Indeed, as expected, the repertoires at 8 days post-infection, when the immune response is strongest (Murali-Krishna et al., 1998),(Slifka et al., 1997) showed evidence of perturbation. Interestingly, LCMV induced a marked decrease in similarity in both V gene and amino acid motif usage in both CD4+ and CD8+ naïve repertoires, perhaps reflecting increased turnover and perturbation of this compartment in response to the infection. However, in contrast to the changes observed in response to chronic environmental antigen stimulation, LCMV drove an increased similarity of effector repertoires. This was reflected not only in V gene and CDR3 nucleotide distributions, but was evidenced by the existence of amino acid triplets highly enriched in the TCR repertoire of infected individuals. Remarkably, many of these triplets were found within the set of CDR3s of CD8+ TCRs which bound one specific epitope of LCMV, confirming the link between motifs and specific antigen recognition. Thus, exposure to a strong synchronous source of antigen, such as is provided by acute exposure to LCMV, drives strong convergent evolution and decreased diversity of the TCR effector repertoire, which relaxes partially towards the uninfected state at 40 days post-infection.

The study we present has a number of limitations. The number of individuals analysed was small, limiting the amount of robust statistical analysis which can be carried out. Thus, many of the conclusions we make are based on statistical trends rather than classical statistical significance thresholds. Furthermore, the analysis of the effects of aging are limited to two time points, and would benefit from extension to very young or very old mice. We also recognize that the functional sub-compartments we define are based on a rather simplistic and limited panel of antibody markers, and that in reality the populations we refer to as naïve, central memory and effector certainly contain further heterogeneity which could be explored further in future studies.

In conclusion, we present a novel approach to the analysis of the TCR repertoire which we use to address the fundamental relationship between stochastic and deterministic processes which drive evolution of the adaptive repertoire. The adaptive immune system shows a remarkable capability to preserve high-order structure, as reflected by conserved frequency distributions of V gene and short amino acid linear motifs, while still allowing enormous diversity at individual sequence level. This high order structure is partially preserved but gradually weakened as the adaptive immune system ages. We speculate that this structure is key to maintaining a robust consistent antigen-specific response across a population in the face of the randomness and heterogeneity imposed by the process of imprecise TCR recombination.

## Materials and methods

### Animals

All experiments except for the LCMV infections were carried out using inbred female Foxp3-GFP (C57BL/6 background) mice sacrificed at three months (young) and one year (adults). All animals were handled according to regulations formulated by The Weizmann Institute’s Animal Care and Use Committee and maintained in a pathogen-free environment.

### LCMV infections

Females C57BL/6 mice at 5 weeks old (Envigo) were in injected with Intraperitoneal with the Armstrong LCMV strain. Mice were collected after 8 or 40 days of infection.

### Sample preparation and T cell isolation

Spleens were dissociated with a syringe plunger and single cell suspensions treated with ammonium-chloride potassium lysis buffer to remove erythrocytes. Bone marrows were extracted from the femur and tibiae of the mice and washed with PBS. Samples were loaded on MACS column (Miltenyi Biotec) and T cells were isolated according to manufacturer’s protocol. Bone marrows cells were purified with CD3+ T isolated kit (CD3ε MicroBead Kit, mouse, 130-094-973, Miltenyi Biotec). Splenic CD4+ and CD8+ cells were purified in two steps: (1) CD4+ positive selection (CD4+ T Cell Isolation Kit, mouse, 130-104-454, Miltenyi) (2) the negative cells fraction were further selected for the CD8+ positive cells (CD8a+ T Cell Isolation Kit, mouse, 130-104-07, Miltenyi Biotec). For the tetramers binding reaction, we pooled splenocytes from previously vaccinated mice (5 mice after 8 days of infection) and purified their T cells using the untouched isolation kit (Pan T Cell Isolation Kit II, mouse, 130-095-130, Miltenyi Biotec).

### Flow cytometry analysis and cells sorting

The following fluorochrome-labeled mouse antibodies were used according to the manufacturers’ protocols: PB or Percp/cy5.5 anti -CD4+, PB or PreCP/cy5.5 anti- CD8+, PE or PE/cy7 anti- CD3+, APC anti-CD62L, Fitc or PE/cy7 anti- CD44 (Biolegend). Cells were sorted on a SORP-FACS-AriaII and analyzed using FACSDiva (BD Biosciences) and FlowJo (Tree Star) software. Sorted cells were centrifuged (450g for 10 minutes) prior to RNA extraction.

### LCMV -tetramers staining and FACS sorting

Three monomers (NIH Tetramer Core Facility) with different LCMV epitopes were used: MHCI- NP396-404(H-2D^b^), MHCI- NP205-212(H-2K^b^), MHCI- GP92-101 (H-2D^b^). Tetramers were constructed by binding Biotinylated monomers with PE/APC – conjugated- streptavidin (according to the NIH protocol). Purified T cells were stained with FITC anti-CD4+ and PB anti-CD8+ and followed by tetramers staining (two tetramers together), for 30 min at room temperature (0.6ug/ml). CD8+ epitope-specific cells were sorted from single- positive gates for one type of tetramer.

### Library preparation for TCR-seq

All libraries in this work were prepared according to the published method(Oakes et al., 2017), with minor adaptations for mice. Briefly, we extracted total RNA from CD4+/CD8+/CD3+ T cells (from spleen or bone marrow) of Foxp3-GFP or C57BL/6 mice using RNeasy Micro Kit (Qiagen) and cleaned from excess DNA with DNAse 1 enzyme (Promega). RNA samples were reverse transcribed to cDNA and an anchor sequence at the variable part of the TCR was added using single strand ligation. Ligation products were amplified by PCR in three reactions, using an extension PCR to add Illumina sequencing primers, indices and adaptors. Our modified protocol for mice included specific primers for the constant region of the TCR α or β chain (“GAGACCGAGGATCTTTTAACTGG”,”GCTTTTGATGGCTCAAACAAGG”, for α and β chain respectively). These primers are used in the reverse transcription (RT) and the first two PCR reactions (PCR1: “CAGCAGGTTCTGGGTTCTGGATG”,” TGGGTGGAGTCACATTTCTCAGATCCT”, for α and β chain respectively). Primers in the second round of the PCR included TCR constant region sequence, together with a six base pair Illumina index for multiplex sequencing, six random base pairs to improve cluster calling at the start of read 1, and the Illumina SP1 sequencing primer (PCR2: “ACACTCTTTCCCTACACGACGCTCTTCCGATCTHNHNNH-index-CAGCAGGTTCTGGGTTCTGGATG”, “ACACTCTTTCCCTACACGACGCTCTTCCGATCTHNHNNH-index-GGTGGGAACACGTTTTTCAGGTCCTC”, for α and β chain respectively). In the third round of the PCR, the primers were the SP1 and SP5 Illumina adaptors (PCR3: “CAAGCAGAAGACGGCATACGAGAT “, “AATGATACGGCGACCACCGAGATCTACACTCTTTCCCTACACGACGCTCTTCC”, forward and revers respectively). All PCR reactions were done using KAPA HiFi high fidelity proof reading polymerase (KAPA Biosystems). Libraries were sequenced using NexsSeq 550 (200 bp forward read, 100 bp reverse) (Illumina).

### Pre-Processing and Error Correction for Raw Reads

Data was processed using an in-house pipeline, coded in R. First, we transfer the UMI sequence from the read2 to read1 sequence. Trimmomatic(Bolger et al., 2014) was used to filter out the raw reads containing bases with Q-value ≤20 and trim reads containing adaptors sequences. The remaining reads were separated according to their barcodes and reads containing the constant region for α or β chain primers sequences were filtered (CAGCAGGTTCTGGGTTCTGGATG/ TGGGTGGAGTCACATTTCTCAGATCCT α and β chain respectively), allowing up to three mismatches. Bowtie 2(Langmead and Salzberg, 2012) (using sensitive local alignment parameters) was used to align the reads to the germline V/J gene segments, as found in IMGT germline. The CDR3 nucleotide sequences were translated to amino-acid sequence in two steps. The N-terminal Cysteine was identified by matching it to the V aligned region. Then the C-terminal Phenylalanine was identified by matching it to the J aligned region. Up to one mismatch was allowed per 18-stretch sequence, ending with the Cys or starting at the Phe. CDR3AA sequences were defined according to IMGT convention. To correct for possible sequence errors, we cluster the sequences UMI’s in two steps; (1) UMI’s with highest frequency grouped within a Levenshtein distance of 1. (2) Out of these sequences, CDR3AA sequences (starting from the most frequent sequence in a group) were clustered using a Hamming distance(Hamming, 1950) threshold of 4. Finally, the UMI of each CDR3 sequence was counted, and UMI count reads with one copy number were filtered out. For the entire analysis, sequences were used only if they were fully annotated (both V and J segments assigned), in-frame (i.e., they encode for a functional peptide without stop codons) and with copy number greater than one. In addition, we removed the invariant α chain of the iNKT CDR3 sequence (“CVVGDRGSALGRLHF”(Greenaway et al., 2013), 0.001% from all sequence in our data).

### Statistical Analysis

All statistical analysis was performed using R Statistical Software. For the pre-processing pipeline we used the “ShortRead” package(Morgan et al., 2009). The package “vegan”(Dixon, 2003) was used to measure the Simpson and Shannon indices(Leinster and Cobbold, 2012),(Mehr et al., 2012). We also used it to compute the Horn similarly index(Greiff et al., 2015),(Venturi et al., 2008) and to project the Nonmetric Multidimensional Scaling(Faith, D. P, Minchin, P. R. and Belbin, 1987). The Horn index relies on both overlap and abundancy of sequences, as evaluated by the unique molecular identifier count (UMI count) (Shugay et al., 2014),(Friedensohn et al., 2017). For the PCA analysis we applied the “factoextra” package(A. Kassambara, 2017) and the “ggplot2” (Wickham H, 2009) was used for generating figures.

## List of Abbreviations

TCR: T cell receptor
TCR-seq: High-throughput sequencing of TCR.
BM: Bonne-marrow
SP: Spleen
MHC: Major histocompatibility complex
CDR3: Complementarity determining region three
CDR3NT/CDR3AA: Nucleotide and amino acid sequences of the CDR3
UMI: Unique molecular identifier
CM: Central memory T cells
N: Naïve T cells
Treg: Regulatory T cells
RT: Reverse transcription
cDNA: Complementary DNA
V, D and J: Variable (V), diversity (D) and joining (J) TCR gene segments
CDR3ntVJ: CDR3NT sequences with V and J gene segments
LCMV: Lymphocytic choriomeningitis virus

## Data availability

All DNA sequences from young and adult mice have been submitted to the Sequence Read Archive under identifier PRJNA771880. **https://www.ncbi.nlm.nih.gov/Traces/study/?acc=PRJNA771880&o=acc_s%3Aa**

## Acknowledgements

This study was initiated and conceived by our friend, mentor and colleague Dr. Nir Friedman (last author). Sadly, Nir died after a long battle with illness without being able to complete the work. We have tried to complete this study in the spirit in which it was undertaken, but we are conscious that we fall far short of the insight and clarity of Nir’s remarkable intellect. We dedicate this study to his memory.

BC was supported by a Weston Visiting Professorship from the Weizmann Institute of Science, and by a grant from the Rosetrees Foundation, UK. NF was supported by the Applebaum Family Foundation,

## Supplementary

**SI Table 1:** UMI count and the cells number of each compartment in young or adult mice. Compartments names are naïve, effector, central memory and Treg (N,E,CM,Treg). The tissues are bone marrow (BM) and spleen (SP). The numbers are extracted after running the TCR sequencing.

|  |  | UMI count |  |  |  |  |  | Cells number |  |  |  |  |  |
| --- | --- | --- | --- | --- | --- | --- | --- | --- | --- | --- | --- | --- | --- |
|  |  | Young |  |  | Adult |  |  | Young |  |  | Adult |  |  |
| Name | Chain | M1 | M2 | M3 | M1 | M2 | M3 | M1 | M2 | M3 | M1 | M2 | M3 |
| CD4+ CM BM | α chain | 976 | 1016 | 3945 | 784 | 22286 | 393 | 6392 | 6000 | 13000 | 8000 | 10000 | 4000 |
| CD4+ CM SP |  | 60059 | 4111 | 69447 | 34621 | 4656 | 7712 | 134768 | 46000 | 88945 | 205000 | 114000 | 51000 |
| CD4+ E BM |  | 12894 | 10729 | 45322 | 10784 | 1354 | 6925 | 66680 | 67000 | 161000 | 70000 | 100000 | 82000 |
| CD4+ E SP |  | 184150 | 68984 | 405212 | 186485 | 161144 | 130397 | 583650 | 226000 | 543631 | 1000000 | 718000 | 650000 |
| CD4+ N BM |  | 4421 | 4101 | 11624 |  |  |  | 26551 | 31000 | 39000 |  |  |  |
| CD4+ N SP |  | 500932 | 241425 | 492564 | 10221 | 9064 | 1711 | 1000000 | 1000000 | 1070000 | 50000 | 58000 | 40000 |
| CD4+ Treg BM |  | 21757 | 7747 | 64357 | 15070 | 9473 | 16730 | 73716 | 99000 | 215000 | 70000 | 50000 | 70000 |
| CD4+ Treg SP |  | 361137 | 164556 | 273854 | 52590 | 54401 | 19406 | 744389 | 400000 | 506092 | 442000 | 162000 | 285000 |
| CD8+ CM BM |  | 12778 | 9895 | 21679 | 7094 | 22467 | 19149 | 28340 | 36000 | 44000 | 47000 | 77000 | 110000 |
| CD8+ CM SP |  | 99513 | 51772 | 159812 | 36879 | 17795 | 40745 | 173090 | 135000 | 178000 | 215000 | 48000 | 160000 |
| CD8+ E BM |  | 7365 | 597 | 3620 | 1684 | 12829 | 81516 | 13667 | 9700 | 22000 | 110000 | 51000 | 800000 |
| CD8+ E SP |  | 10599 | 3609 | 22482 | 15442 | 68036 | 11924 | 33298 | 31000 | 47000 | 120000 | 216000 | 105000 |
| CD8+ N BM |  | 13874 | 4684 | 60752 |  |  |  | 45376 | 112000 | 148000 |  |  |  |
| CD8+ N SP |  | 192821 | 464157 | 596538 | 36490 | 11830 | 18848 | 1000000 | 1200000 | 1200000 | 219000 | 49000 | 73000 |
| CD4+ CM BM | β chain | 1337 | 980 | 2666 | 885 | 1834 | 539 | 6392 | 6000 | 13000 | 8000 | 10000 | 4000 |
| CD4+ CM SP |  | 62857 | 3235 | 60734 | 48330 | 5172 | 8143 | 134768 | 46000 | 88945 | 205000 | 114000 | 51000 |
| CD4+ E BM |  | 32517 | 25937 | 47924 | 14497 | 41405 | 9936 | 66680 | 67000 | 161000 | 70000 | 100000 | 82000 |
| CD4+ E SP |  | 219390 | 85408 | 315338 | 263168 | 174077 | 101827 | 583650 | 226000 | 543631 | 1000000 | 718000 | 650000 |
| CD4+ N BM |  | 4700 | 5024 | 13424 |  |  |  | 26551 | 31000 | 39000 |  |  |  |
| CD4+ N SP |  | 642033 | 197314 | 415101 | 16344 | 11544 | 2364 | 1000000 | 1000000 | 1070000 | 50000 | 58000 | 40000 |
| CD4+ Treg BM |  | 19707 | 8123 | 60971 | 14335 | 14906 | 12208 | 73716 | 99000 | 215000 | 70000 | 50000 | 70000 |
| CD4+ Treg SP |  | 455374 | 199681 | 234393 | 157530 | 48780 | 21502 | 744389 | 400000 | 506092 | 442000 | 162000 | 285000 |
| CD8+ CM BM |  | 13442 | 9438 | 19861 | 9031 | 34436 | 14826 | 28340 | 36000 | 44000 | 47000 | 77000 | 110000 |
| CD8+ CM SP |  | 104644 | 56797 | 145804 | 38910 | 19600 | 61581 | 173090 | 135000 | 178000 | 215000 | 48000 | 160000 |
| CD8+ E BM |  | 5229 | 929 | 3945 | 2414 | 18903 | 132048 | 13667 | 9700 | 22000 | 110000 | 51000 | 800000 |
| CD8+ E SP |  | 11077 | 3636 | 15769 | 23605 | 90234 | 14581 | 33298 | 31000 | 47000 | 120000 | 216000 | 105000 |
| CD8+ N BM |  | 16392 | 6490 | 55300 |  |  |  | 45376 | 112000 | 148000 |  |  |  |
| CD8+ N SP |  | 231312 | 487456 | 515372 | 53474 | 17830 | 14301 | 1000000 | 1200000 | 1200000 | 219000 | 49000 | 73000 |

**SI Figure 1:**
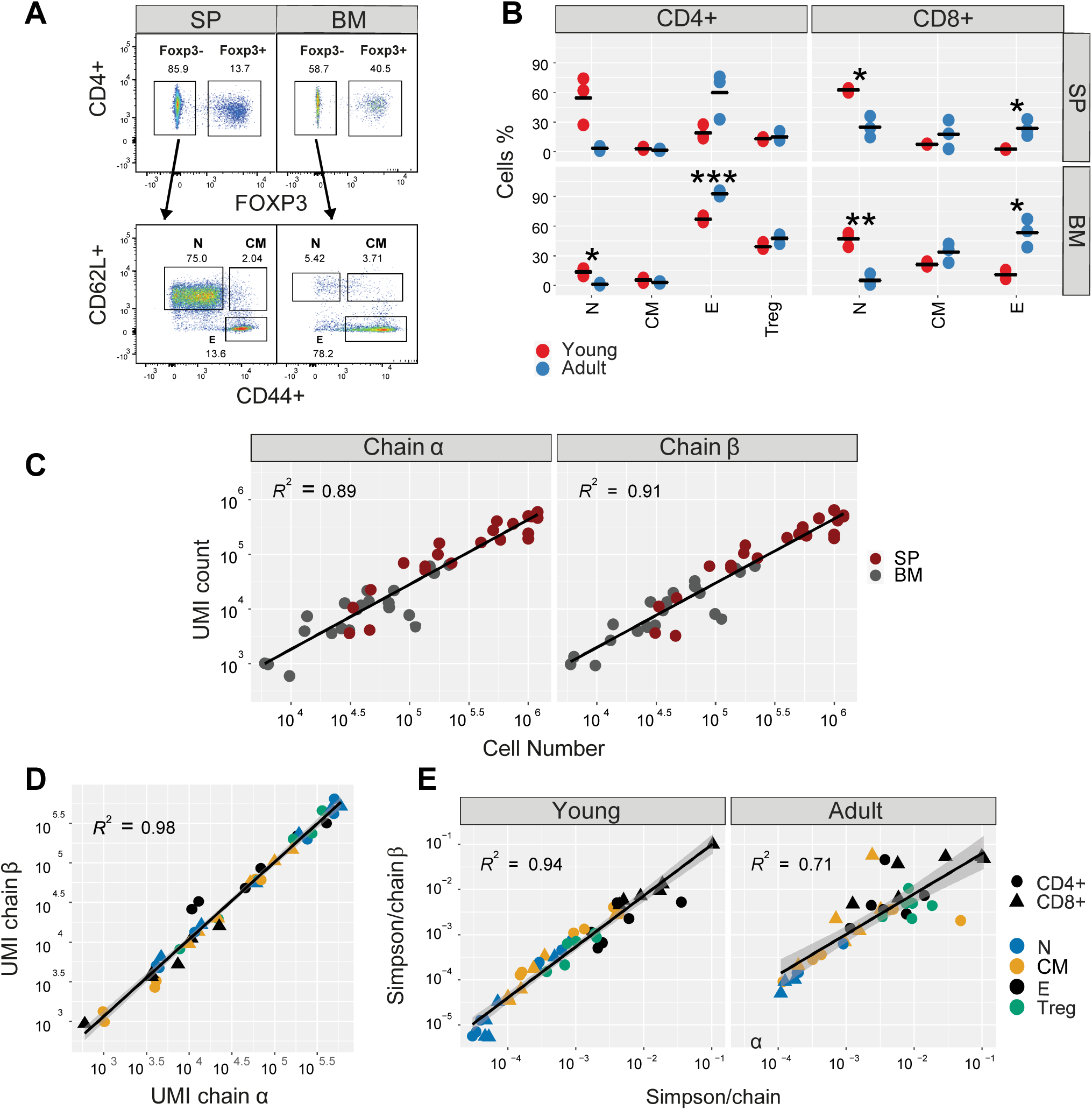
(A) Representative sorting gates for CD4+ cells of one young mouse. (B) FACS-sorting cells percentage of each compartment of young (red) or adult (blue). Mean is shown in black lines (n=3). Significant differences between age groups are denoted by asterisks (P-values: ^*^ <0.05, ^**^ < 0.01, ^***^ <0.001, t-test). (C) The number of obtained UM correlates with sorted cells number. Colored dots correspond to the sum of UMI count in the shown young mice compartments vs. the number of sorted cells. α and β chains are marked in circles and triangles, respectively. P-value= 1.83×10-42, R^2^ = 0.9. (D) High correlation between α and β UMI counts. Colored dots correspond to the sum of each young mice compartment (color and shape). (E) Shannon indices from α and β repertoires are highly correlated. Each point is the Shannon index of one SP or BM, CD4+ or CD8+ (dots shape) compartments from young or adult mice (upper or lower panel, respectively).

**SI Figure 2:**
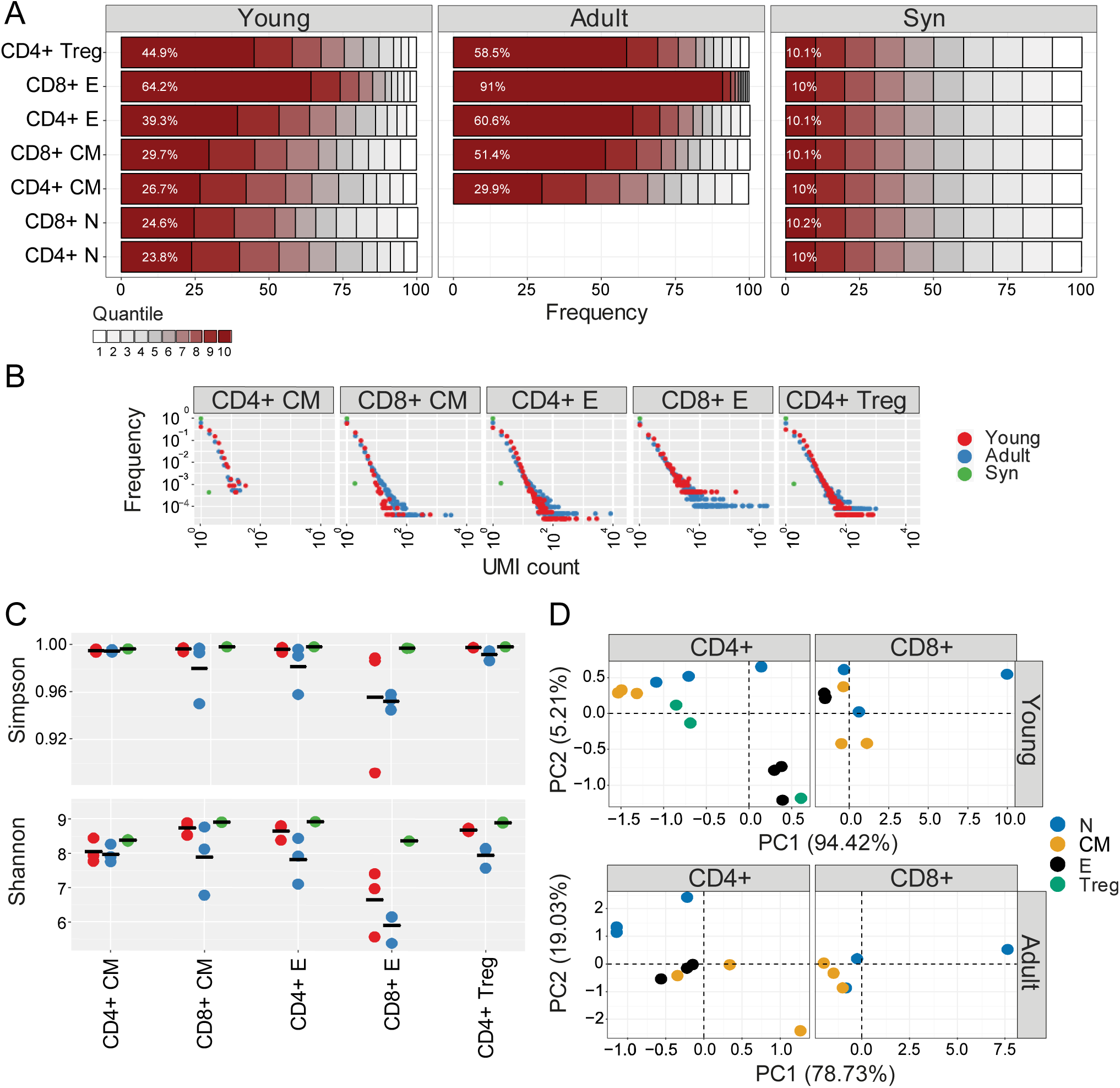
Clonal expansion and diversity of the TCRβ repertoire in different bone marrow subsets of young and adult mice. (A) The TCRs in each repertoire were ranked according to frequency. The proportion within each decile is illustrated (low abundance sequences in white, ranging to high abundance sequences in dark red). The percentage of the distribution represented by the top decile is shown in white text. (B) The sequence abundance distribution in each compartment. The plots show the proportion of the repertoire (y-axis) made up of TCR sequences observed once, twice, etc. (x-axis). Repertoires from young mice are shown with red dots, older mice with blue dots, and synthetic repertories in green. (C) Simpson and Shannon scores of equal repertories size (500 CDR3NT’s) from each compartment and mouse. Colors same as panel B. Mean is shown in black lines (n=3). (D) PCA of the Renyi diversities of order 0, 0.25, 0.5, 1, 2, 4. CD4+ or CD8+ T cells compartments (color dots) from young or adult (left or right panel respectively).

**SI Figure 3:**
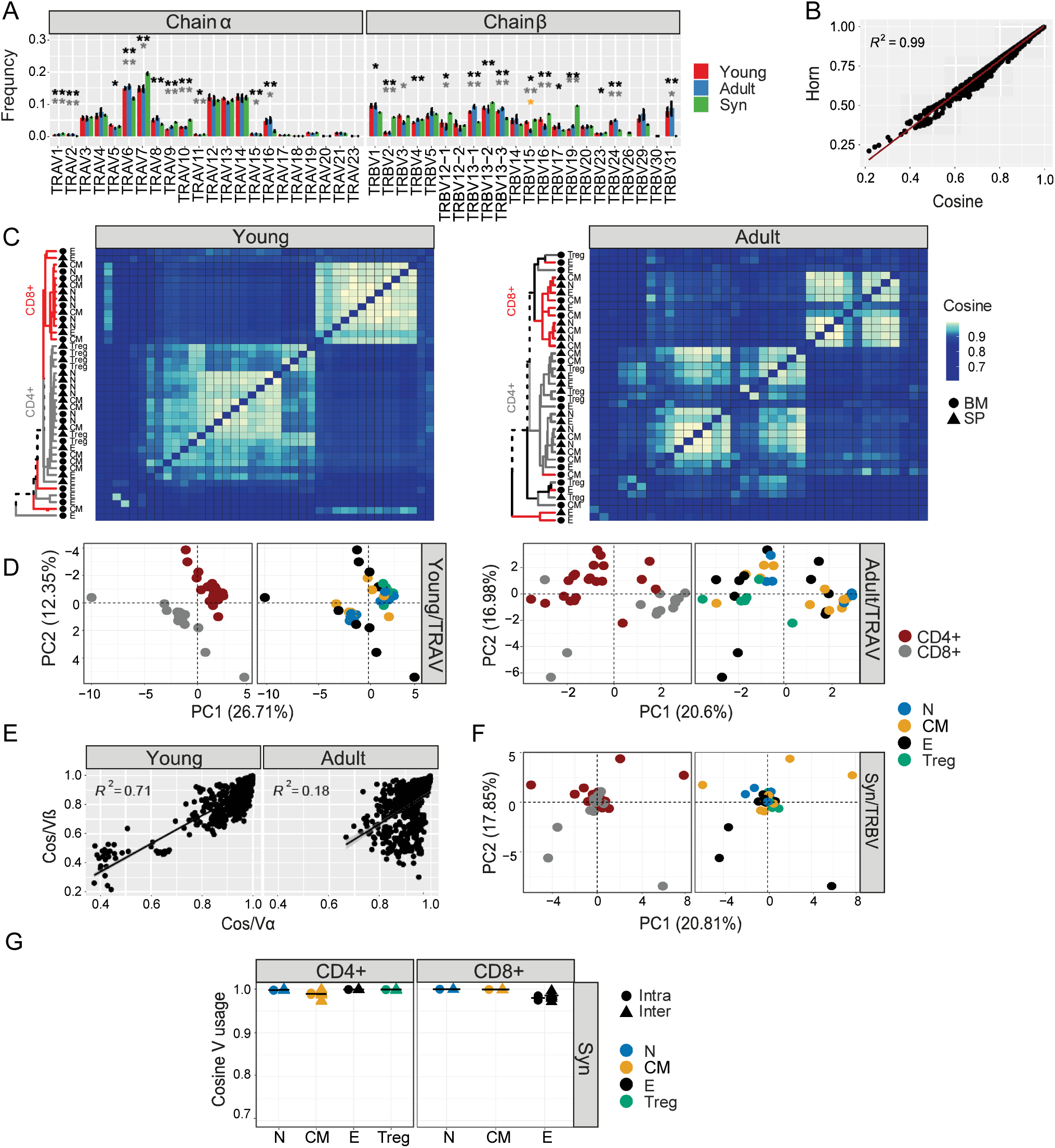
(A) TRV usage of naïve cells from young (red), adult (blue), and synthetic (green) mice. Each bar represents the mean frequency of the V segment in grouped naïve T cells from both tissues. Error bars are SEM (n=6, three mice from CD4+ and CD8+ naïve). Significant differences between all pair groups (Young vs Adult= orange, Young vs Syn=black, Adult vs Syn=grey) in specific segments are detected both in TRBV genes and TRAV families of genes (P-values: ^*^ <0.05, ^**^ < 0.01, t-test with Benjamini & Hochberg correction). (B) A high correlation between Cosine and Horn similarity measurements was calculated for the TRBV usage. Each point is the Horn or Cosine score for the Vβ usage between all pair compartments. (C) The cosine similarity index of the TRAV usage was calculated between all pairs of repertories in young (left) or adult (right) mice. Hierarchical clustering dendrograms show the organization of the assigned at each plot, colored by CD4+ and CD8+ groups (grey and red branches respectively) and labels by compartment (text and symbol). Tissues are marked in symbols shape (SP= triangles, BM= circles). (D) PCA separates the Vα usage between CD4+ and CD8+ class of young (upper) or adult (lower) mice but not within their subgroup compartments. Each color represents one compartment from one mouse (e.g., CD8+ Effectors, BM, mouse 1). (E) Pairwise cosine similarities between Vα and Vβ usage show low correlation, especially in adult mice. Each point is the cosine similarity for Vα and the Vβ usage. (F-G) Uniform Vβ usage in synthetic TCRs, both in PCA analysis (F) and in pairwise cosine similarity scores (G).

**SI Figure 4:**
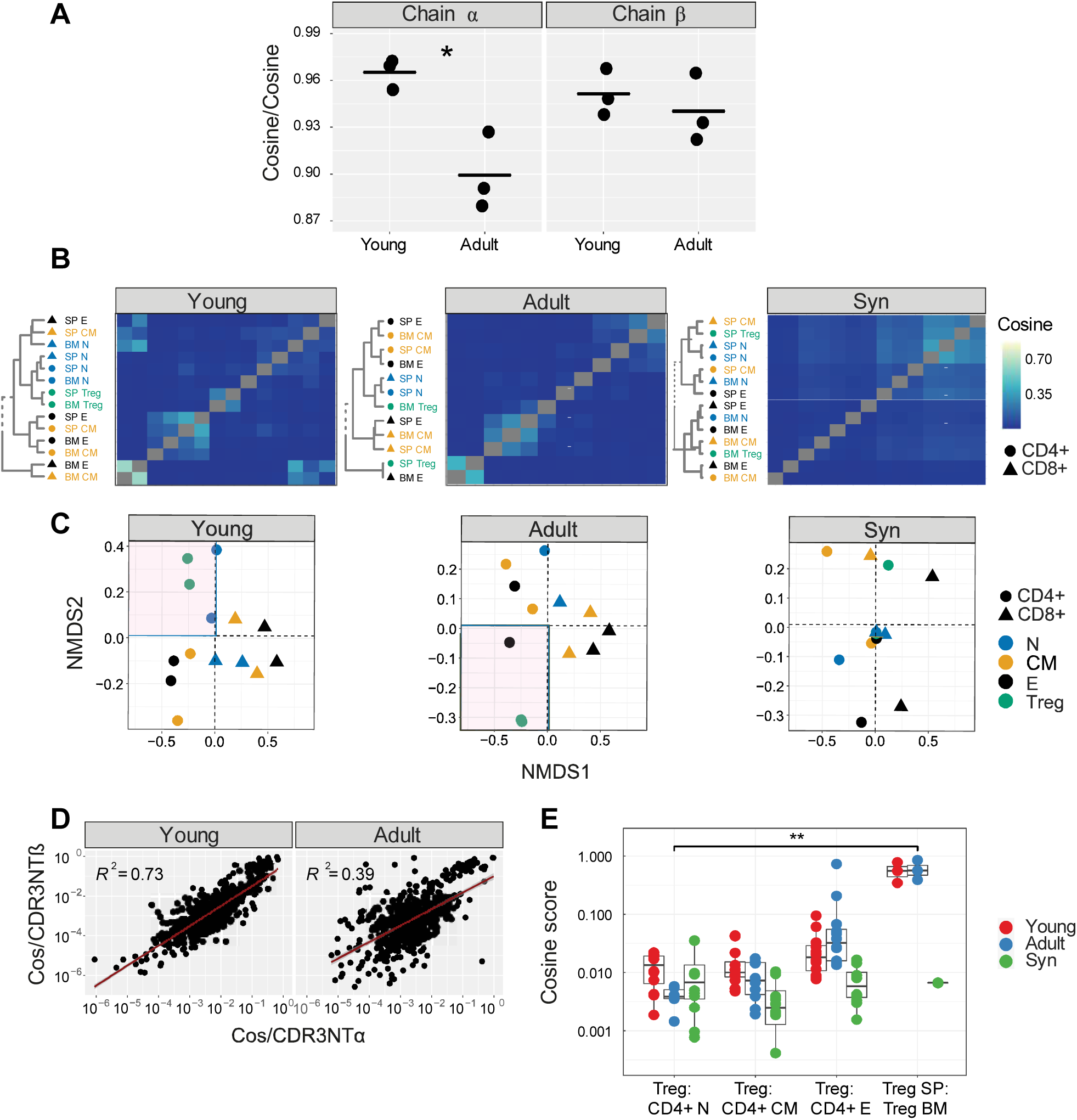
Differential sharing of T cell CDR3 nucleotide α and β chain sequences defines different subpopulations of T cells. (A) Similar CDR3NT α and β Cosine scores across young and adult mice. Cosine measurement calculated for CDR3NT between all pair compartments within each young or adult mouse (for example, in young mouse 1: Treg SP and CD4+ N BM). These values were compared across mice using another Cosine score calculation. The dots color corresponds to the TCR chain (red= TCRα, grey= TCRβ). Significant differences between age groups are denoted in asterisks (P-values: ^*^ <0.05, ^**^ < 0.01, t-test). (B) Pairwise cosine similarity from representative young, adult, or synthetic (“Syn”) mouse CDR3αNT sequences. Correlation levels are represented by color (high=light blue, low= dark blue). In color and text, hierarchical clustering dendrograms for all T cell compartments are plotted to the left of each heat map (CD4+=circle, CD8+= triangles). (C) The similarity matrices shown as heatmaps in B are represented in two dimensions by NMDS. (D) CDR3αNT vs. CDR3βNT pairwise cosine similarities between all pairwise compartments of young and adult mice. (E) Cosine index sharing levels between CDR3βNT of Tregs across tissues or naïve and CD4+ effector repertoires within each young(red), adult(blue) or synthetic-based (green) mouse. Comparisons between the different tissues (SP-SP, SP-BM, BM-BM, n= 9). Mean is shown by horizontal black lines. Significant differences are denoted in asterisks (P-values: * <0.05, **< 0.01, T-test) and calculated between the groups: Tregs across tissues and Treg CD4+ naïve cells.

**SI Figure 5:**
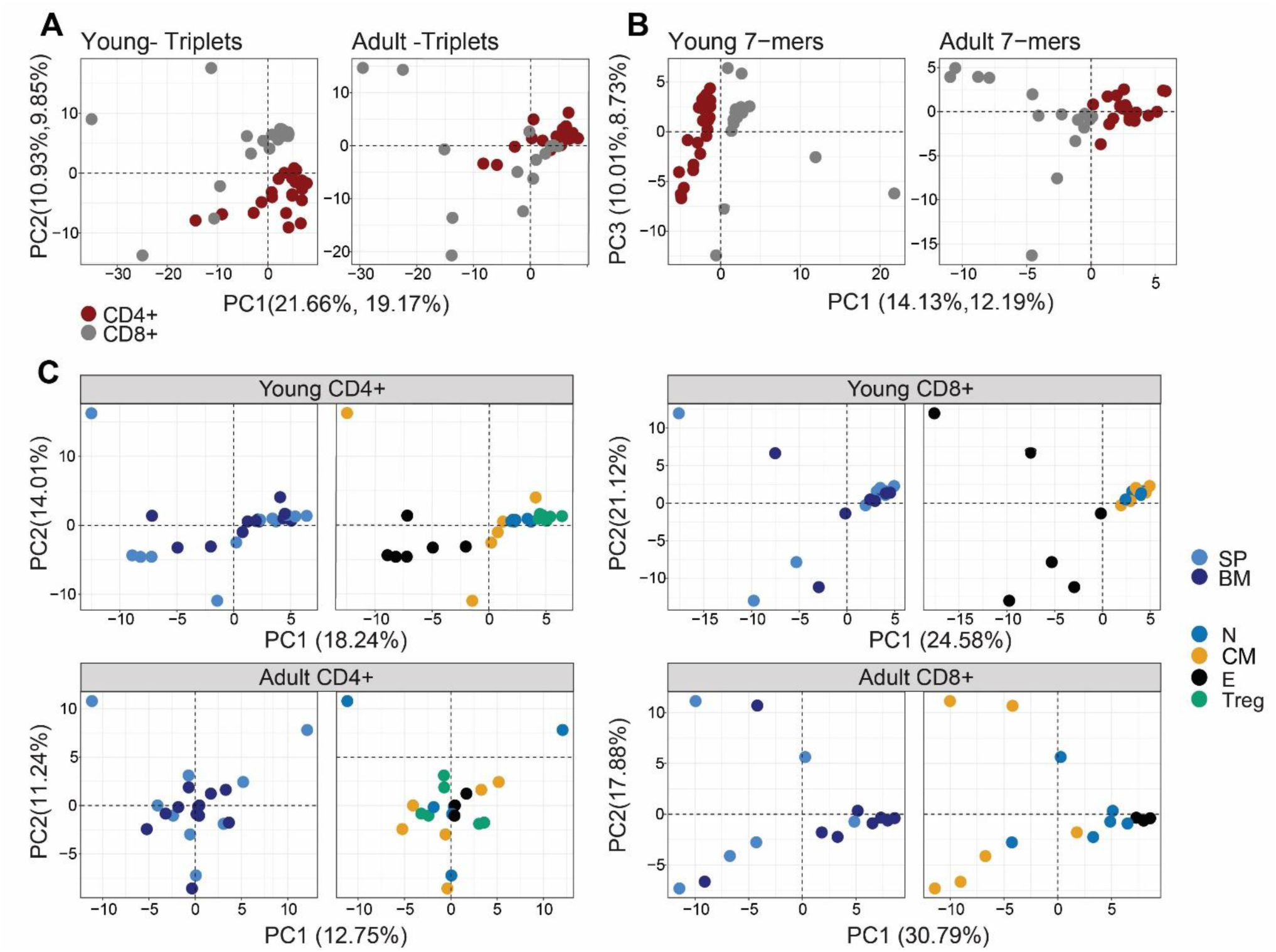
CD4+ T cells compartments distinct top CDR3βAA motifs, alter with age. Top triplets and 7-mers are selected by the mean frequency of each motif across all compartments and mice (A-B) PCA analysis of the top CDR3AAβ triplets (A), and 7-mers (B) motifs separate between CD4+ and CD8+ class (red and grey dots, respectively) in young (left) and adult (right) mice. (C) CDR3βAA 7-mers PCA analysis of CD4+ (left) or CD8+ (right) from young (upper) or adult (lower) mice. See legend for symbols and color code.

**SI Figure 6:**
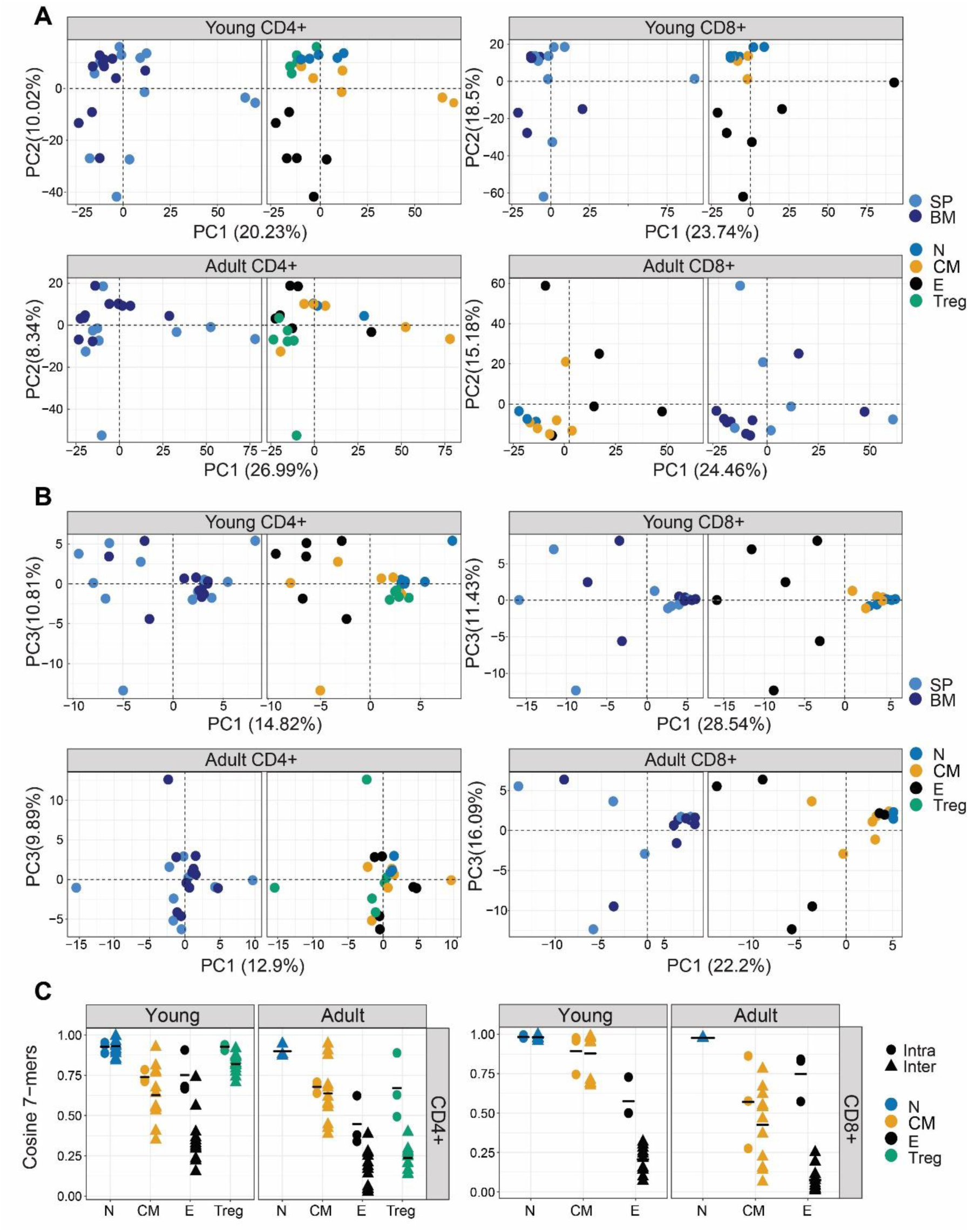
PCA analysis of the top CDR3αAA motifs separates between CD4+ compartments of young mice, yet in a slightly lower degree than the CDR3βAA motifs. (A-B) PCA analysis of the top CDR3αAA triplets (A) or 7-mers motifs (B). CD4+ or CD8+ compartments (left and right, respectively) in young or adult mice (upper and lower, respectively) are assigned in color dots. (C) Pairwise cosine similarities scores of the top 7-mets CDR3βAA motifs between individuals (circles) or within individuals (between spleen and bone marrow, triangles). T cells compartments (colored dots) are divided into CD4+ (left) and CD8+ (right) from young or adult mice. Mean is shown by horizontal black lines.

**SI Figure 7:**
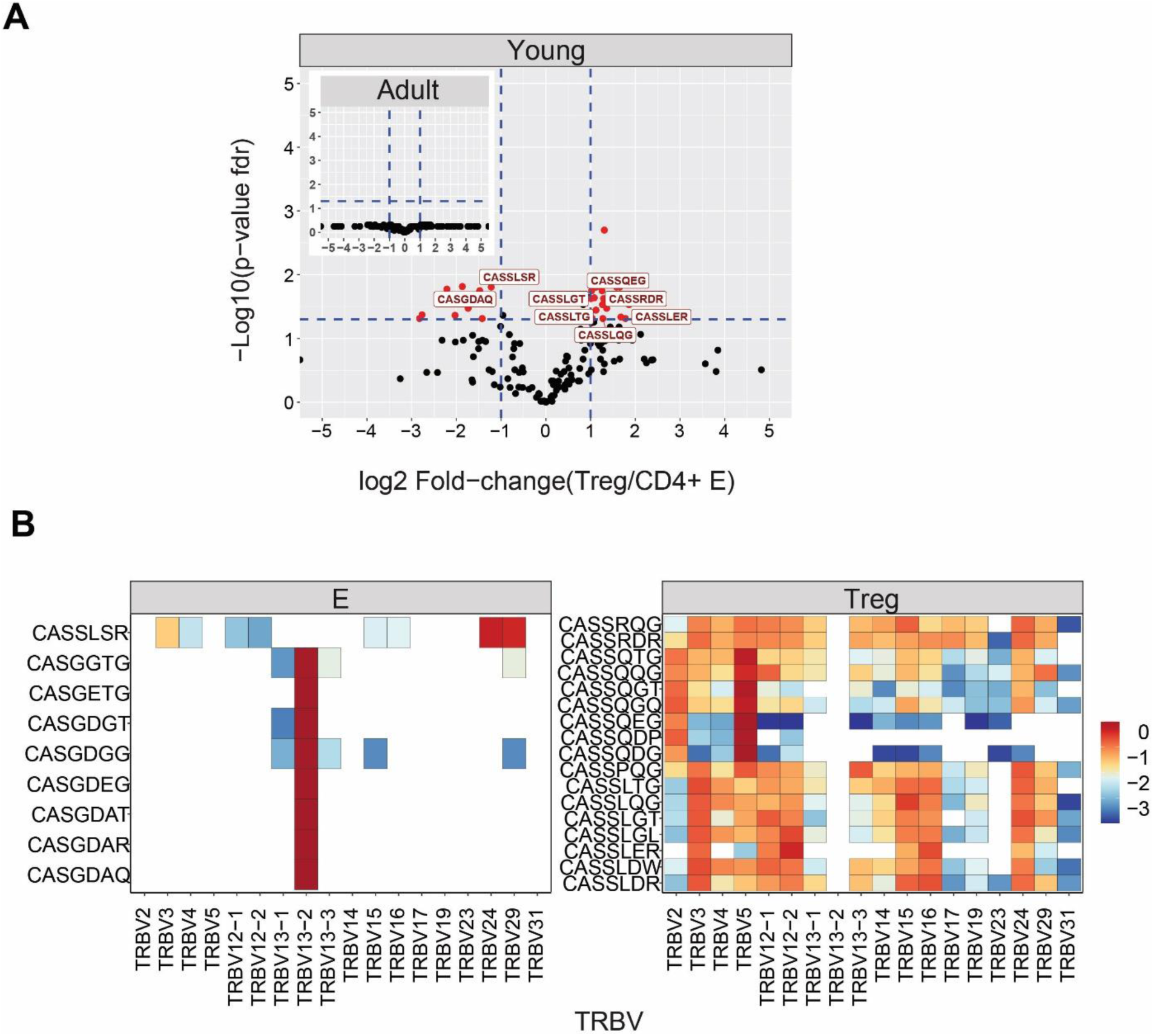
CD4+ T cell compartments express distinct 7-mers β chain motifs in young and not adult mice. (A) Treg and CD4+ effector differentially expressed 7-mers are found in young but not adult mice. Each dot represents a single 7-mer motif. P-value (t-test) was calculated for each motif across six samples (three mice and two tissues) of CD4+ Treg and CD4+ effector cells. The Y-axis shows FDR-adjusted p-values. The X-axis shows the log 2-fold-change, calculated between Treg and CD4+ effector mean motifs frequency across compartments (6 samples each). Significance thresholds are marked in blue lines: (1) at y=1.3 (equivalent to a p-value of 0.05) and x=±1 (denoting a total fold-change of 2). Representative 7-mers above both thresholds are labeled with red text and dots. (B) The Vβ usage of the CD4+Treg (right) and CD4+ effector (left) differentially expressed 7-mers. The color represents the log10 frequency of each 7-mer in a specific Vβ gene (low= blue, high=red).

